# ORB-TXTL: cell-free expression of membrane proteins on lipid bilayer-coated beads

**DOI:** 10.64898/2026.08.26.747166

**Authors:** Aset Khakimzhan, Seth Thompson, Vincent Noireaux

**Affiliations:** School of Physics and Astronomy, University of Minnesota, Minneapolis, MN 55455, USA

## Abstract

Membrane proteins achieve a remarkable range of cellular functions, yet their characterization at high throughputs remains difficult with standard reconstitution methods. Here, we develop On-bead Reconstitution into Bilayers via Cell-free Transcription and Translation (ORB-TXTL), a platform that uses compositionally tunable lipid bilayer-coated silica beads as scaffolds for cell-free synthesized interacting and integral membrane proteins. ORB-TXTL is fast as it just takes a few hours to integrate membrane proteins onto the beads, which can be extensively washed and seamlessly transferred between reaction buffers, to perform assays that are read out by standard laboratory equipment without tagging and sophisticated equipment. We first characterized the lipid interactions of the mechanosensitive channel MscL, then screened 169 *E. coli* proteins and identified a systematic dependence of membrane integration efficiency on the number of transmembrane domains. Finally, we functionally reconstituted the *E. coli* phospholipid synthesis pathway, demonstrating that ORB-TXTL is a tractable and cheap chassis for multi-enzyme membrane biochemistry.

## Main

With advances in computational protein structure prediction and generation, platforms for rapid synthesis and characterization of proteins are becoming increasingly valuable. For interacting and integral membrane proteins (IIMPs), which achieve a broad range of essential functions in living cells^1^, such workflows often require tailored synthesis and isolation protocols for each protein of interest and become cumbersome at higher throughputs. Cell-free gene expression (CFE) is an alternative approach to IIMPs synthesis that circumvents some of these issues by directly producing proteins of interest in lipidic membrane environments^2–4^.

IIMPs synthesized by CFE (also known as cell-free transcription-translation, TXTL) are usually reconstituted in liposomes^5,6^, extracellular vesicles^7^, nanodiscs^8–10^, lipid nanoparticles^11^, or using detergents^12–14^ . Each of these methods has enabled advances in IIMPs characterization and synthetic cell research^15^. However, these platforms have technical limitations, such as not supporting repeated washing steps, rapid buffer exchanges, and automated liquid handling, which are all beneficial for increasing throughput and reducing experimental variability. Previous foundational work introduced cell-free IIMPs synthesis and integration into supported lipid bilayers coated silica beads^16,17^. In these pioneering studies, the manipulability of the beads, the control over the lipid bilayer composition, and the high-throughput capacity of this system were not thoroughly explored.

Here, we introduce On-bead Reconstitution into Bilayers via Cell-free Transcription and Translation (ORB-TXTL). This technique simplifies the fabrication of lipid bilayer-coated silica beads as a scaffold for IIMPs synthesis and is highly effective for pairing protein-lipid interaction screening with IIMPs activity assays. The first advantage of ORB-TXTL is the ease of bead manipulation via common pipettors, which enables extensive washing procedures using standard benchtop spinners and multistep reactions with minimal sample loss. Second, the silica surface of the beads lends itself to rapidly assembling stable and compositionally tunable SLBs. And third, ORB-TXTL is compatible with standard molecular biology readouts such as fluorescence, protein electrophoresis, thin-layer chromatography (TLC), and microscopy – making the tool accessible to a broad community of scientists interested in IIMPs.

In this work, we leveraged our previous work with silica sensors^18^ to rapidly fabricate SLBs on 5 μm silica beads. The protein-lipid interaction of the mechanosensitive protein MscL observed with the beads was consistent with the label-free measurements with the resonating sensors, and established ORB-TXTL as an accessible and rapid platform to faithfully characterize protein-lipid interactions. We utilized the speed advantage of ORB-TXTL to conduct a large-scale protein-lipid interaction screen of 169 *E. coli* proteins, identifying a clear dependence of spontaneous membrane integration efficiency on the number of transmembrane domains. Finally, we applied ORB-TXTL to functionally reconstitute the *E. coli* phospholipid synthesis pathway, establishing the platform as a chassis for multi-enzyme membrane biochemistry.

## Results

### Formation of lipid bilayers on beads enables cell-free membrane protein expression

The fabrication of SLBs on silica beads was achieved in about one hour by the following protocol, modified from the solvent-assisted lipid bilayer (SALB) formation method: 5 μm silica beads were incubated in isopropanol, then gently mixed in a lipid-isopropanol solution for 25 minutes; afterwards, SLBs were formed by sequential replacement of the lipid-isopropanol solution with saline buffer^19^ (10 mM Tris, 150 mM NaCl, pH 7.5) (**Figure 1a**). When lipids were not supplied to the solvent, an SLB did not form, while in cases where either pure DOPC or a complex *E. coli* lipid (ECL) mixture was provided, the SLBs formed and were visible when stained with the lipophilic dye Nile Red (**Figure 1b**). Compared to DOPC (1 mg/ml), ECL required greater lipid concentrations (3 mg/ml) to reach saturation coverage (**Figures S1, S2, S3**), which is consistent with QCM SLB fabrication data (**Figure S4**).

**Figure 1.**
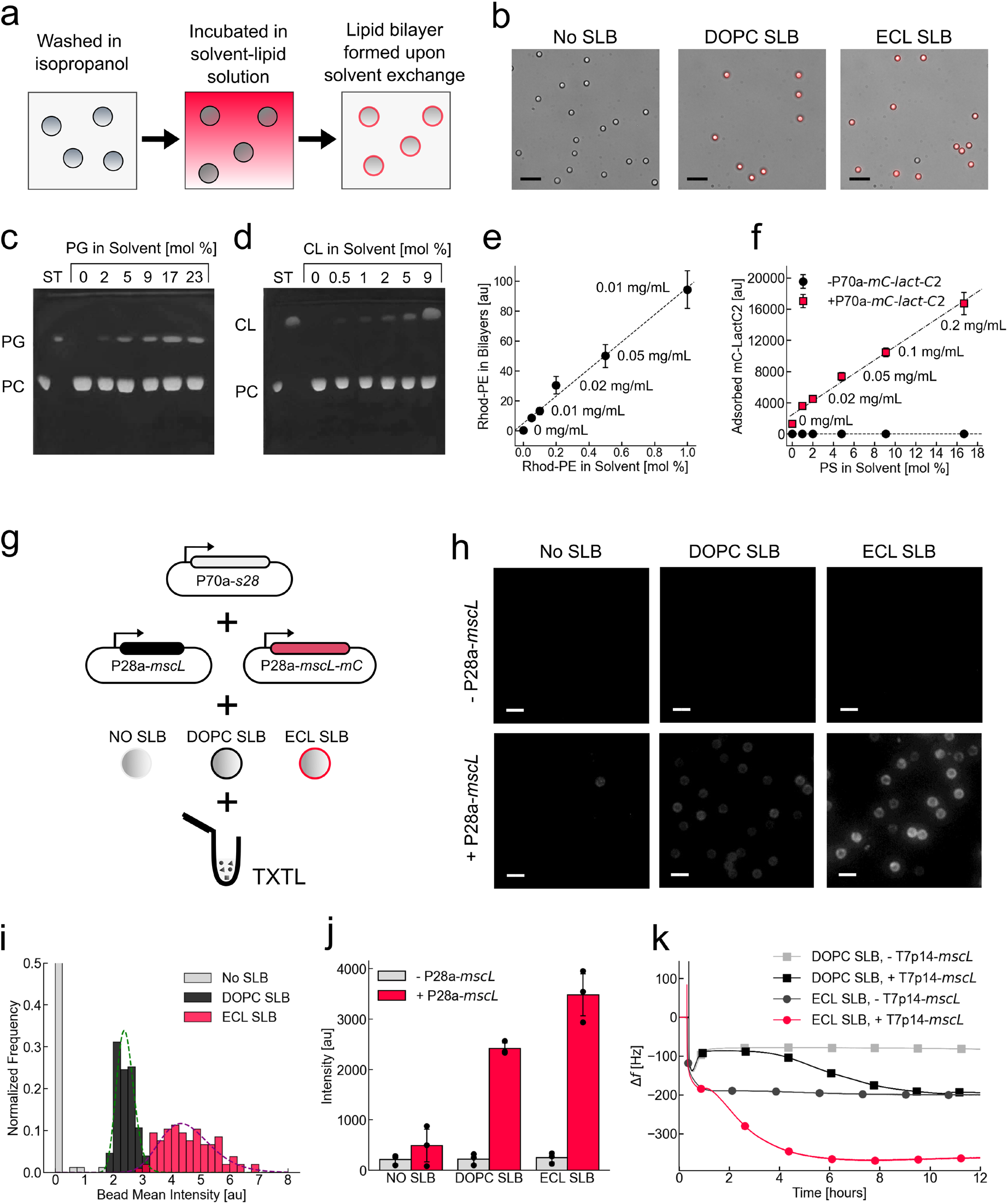
Formation of SLBs on silica beads for IIMPs cell-free synthesis and purification. **(a)** A schematic illustrating the process of forming SLBs on the surface of solid silica beads. **(b)** Images of different SLB compositions on silica beads treated with Nile Red. 20 µm scale bar. **(c) and (d)** Thin-layer chromatography images from silica beads with increasing quantities of PG and CL in the lipid-solvent solution added to DOPC. **(e)** Mean intensity of lipid bilayers with increasing quantities of Rhodamine-PE in the lipid-solvent solution. **(f)** Measuring the binding of mC-Lact-C2 to lipid bilayers with increasing concentration of PS. **(g)** A schematic for assembling an ORB-TXTL reaction that expresses mscL and mCherry-tagged mscL on silica beads with either DOPC or ECL lipid bilayers. **(h)** Red-channel fluorescence images of the experiments outlined in **(g)**, 10 µm scale bar. **(i)** Distribution of the mean red-channel fluorescence intensities for beads with DOPC or ECL SLBs interfaced with TXTL expressing *mscL* and *mscL-mC*. **(j)** A plate-reader fluorescence measurement of the experiments outlined in the prior histogram. **(k)** A QCM-TXTL run demonstrates how MscL is synthesized and integrated in real-time with planar SLBs on planar QCM sensor silica surfaces.

While highly anionic lipids such as PG (phosphatidylglycerol), CL (cardiolipin), and PS (phosphatidylserine) could not form stable homogeneous mono-lipid SLBs on silica due to electrostatic repulsion at physiological pH^20^, they can be incorporated in controlled proportions to other SLBs. On DOPC-based SLBs on beads, PG and CL were added at up to 23% and 9% (molar), respectively, without affecting total coverage (**Figures 1c and 1d**). The addition of Rhodamine-PE into DOPC-based SLBs confirmed that lipid addition is near-linear (**Figure S5, Figure 1e**), which is a valuable feature for quantitative assessment of protein-lipid interactions. To demonstrate the point, we measured the binding of the TXTL-synthesized fusion protein mC-Lact-C2 (a fusion of the fluorescent reporter protein mCherry (mC) and the PS-binding Lact-C2^21^), and observed the binding to also be near-linear relative to the concentration of PS in the SLBs (**Figure 1f**).

To characterize membrane protein integration into silica bead SLBs, we synthesized the well-studied mechanosensitive channel and widely used model membrane protein *E. coli* MscL^22^. Each reaction contained 15 μL of TXTL solution and 2 μL of concentrated bead suspension, corresponding to 63 nL of bead solid volume and 0.75 cm² of surface area. Constructs encoding untagged MscL (P28-*mscL*) and mCherry-tagged MscL (P28a-*mscL-mC*) were co-expressed in the presence of beads under three lipid conditions (No SLB, DOPC, ECL) (**Figure 1g**). The optimal ratio of untagged to tagged DNA was empirically determined as 8:1, as it produced the strongest signal (**Figure S6**). That optimum ratio is consistent with the pentameric stoichiometry of MscL, where such a ratio would favor complexes with a single fluorescent subunit.

Following a 12-hour incubation with the TXTL reaction, beads were washed in a Hepes buffer matching the salt composition of TXTL (50 mM Hepes pH 7.5, 150 mM K-glutamate, 10 mM Mg-glutamate). As an *E. coli* integral membrane protein, MscL integrated more strongly into ECL than DOPC SLBs, observed by the greater fluorescence intensity on ECL than on DOPC SLBs (**Figure S7, Figure 1h**). The population statistics of the fluorescence of isolated beads were quantified to determine both the difference in membrane protein association with a given SLB lipid composition and its reproducibility — DOPC SLBs showed lower mean intensity but lesser bead-to-bead variability (CV of 12% and 19%) than ECL, which we attribute to fluctuations caused by the compositional complexity of the *E. coli* lipid mixture (**Figure 1i**). Across three independent biological replicates, plate reader measurements recapitulated these results with high reproducibility (CV of 4% and 12% for DOPC and ECL, respectively; **Figure 1j**). The consistent, scalable, and rapid ORB-TXTL readouts are enabled by the ease of washing and centrifuging beads, which are not accessible to other approaches. For example, liposome-based systems would require time-consuming high-speed centrifugation steps to minimize liposome sample attrition and nanodisc-based workflows would require size exclusion or affinity chromatography to obtain equivalent samples.

We compared the MscL interaction strength on beads with results using a Quartz Crystal Microbalance (QCM), which can be applied as a label-free instrument to measure IIMPs integration kinetics into supported lipid bilayers fabricated from similar lipid compositions^18^. The QCM sensors were also composed of silica, thus offering the same surface chemistry for both approaches and making them electrostatically equivalent to a nascent protein. A decrease in the QCM sensor resonance frequency corresponds to tightly adsorbed mass on the sensor surface, and the rate of this shift can be used to infer how rapidly proteins were synthesized into the bilayer. The QCM-based ECL SLBs showed a larger frequency drop (Δf ≈ −150 Hz) that saturated in 6 hours, compared to DOPC SLBs (Δf ≈ −100 Hz) that saturated over 12 hours (**Figure 1k**), mirroring the greater fluorescence intensity of ECL-covered beads compared to DOPC-covered beads. No frequency drop was observed in the absence of MscL-encoding DNA, confirming that the signal reflects membrane integration of synthesized protein rather than nonspecific adsorption.

### Large-scale targeted screening reveals patterns of cell-free synthesized membrane protein interaction and insertion

We applied ORB-TXTL to 169 *E. coli* proteins spanning a range of membrane topologies and functional classes, revealing novel protein-lipid interactions and systematic protein integration patterns. The DNA library was prepared by a two-step PCR directly from *E. coli* K12 genomic DNA, using universal primers containing the *E. coli* constitutively expressed P70a promoter and T500 terminator, eliminating the need for plasmid-cloned constructs (**Figure 2a, Figure S8**). Each gene was then expressed for 12 hours in four conditions in 96-well plates: no beads, bare silica beads (No SLB), DOPC SLBs, and ECL SLBs. After the reaction was complete, the beads were washed in Hepes buffer and eluted in 10 µL of SDS-PAGE sample buffer for analysis by SDS-PAGE.

**Figure 2.**
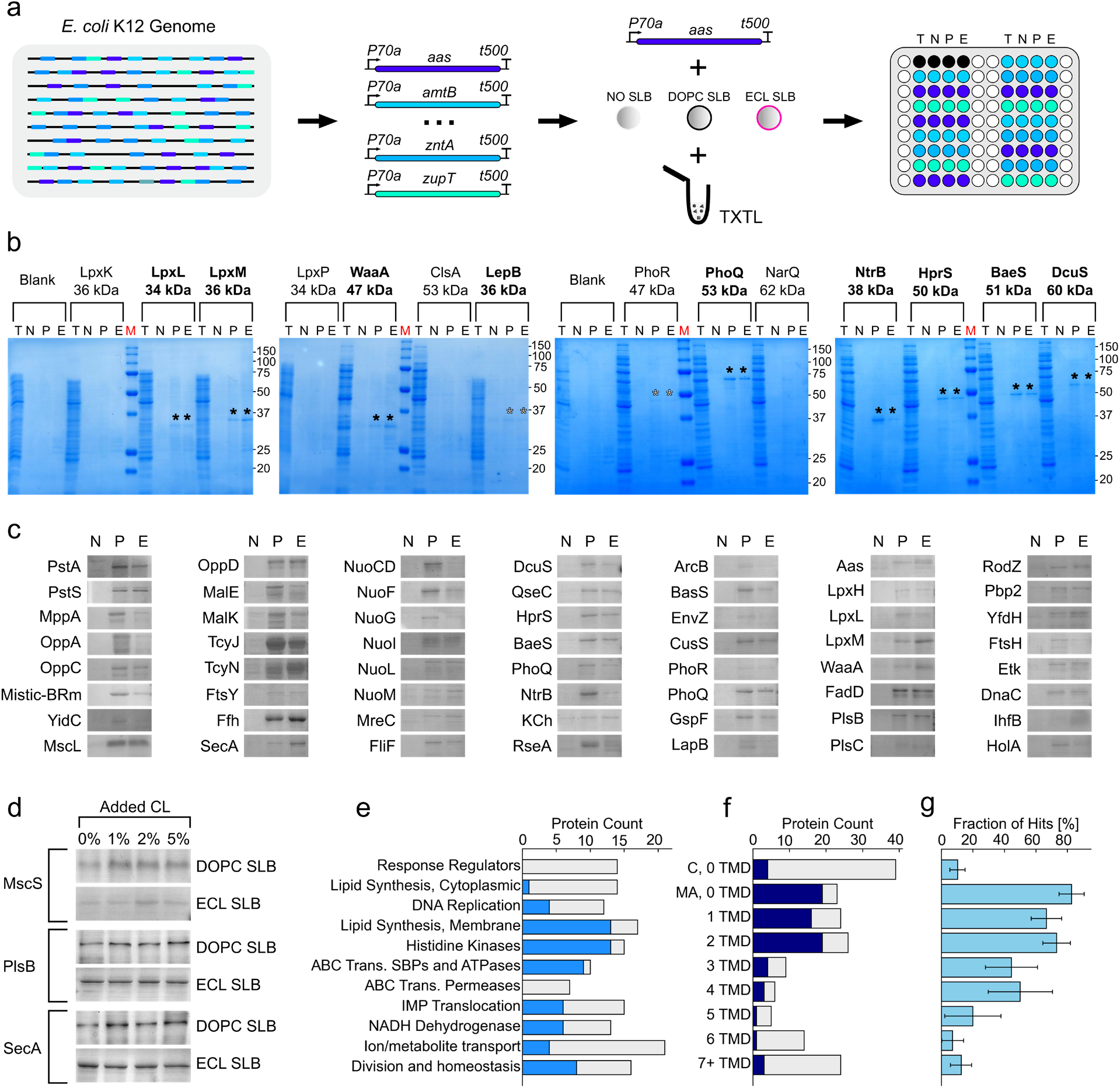
Large-scale protein-lipid interaction screens using SLB-coated beads. **(a)** A schematic for preparing candidates for the interaction campaign. First, genes were extracted from *E. coli* K12 and functionalized with the P70a promoter and T500 terminator. Second, each gene was added to TXTL reactions that contain either no beads (T), beads with no SLB (N), beads with DOPC SLB (P), or beads with ECL SLBs (E). The reactions were parallelized and incubated in a 96-well plate for 12 hours, after which they could be assayed or analyzed for SLB association using SDS-PAGE. **(b)** Examples of SDS-PAGE from such an experiment, with hits observed for both lipid A synthesis enzymes and a selection of Histidine Kinases. The letters ‘TNPE’ above the gels indicate which bead condition is paired with the reaction, while the red M indicates ladder lanes. Filled stars above the bands were placed above intense bands, and hollow stars were placed above bands that were difficult to detect. **(c)** Cropped gel sections to summarize the screen protein hits. **(d)** Dependence of MscS, PlsB, and SecA interactions on the concentration of CL in DOPC and ECL SLBs. **(e)** A total count of protein hits depending on their cellular role and predicted location. **(f)** A total count of protein hits depending on the predicted number of TMDs. The ‘C, 0 TMD’ condition stands for cytosolic proteins, while ‘MA, 0 TMD’ stands for membrane-associated proteins with zero predicted TMDs. **(g)** The rate of protein hits depends on the number of predicted TMDs.

The screen produced interpretable gels across all 169 genes, with bands scored as hits when the signal exceeded five times the standard deviation of the local background (**Figure 2b, Figures S9-24**). The results are organized in a summary file (Supplementary Data 1). For most hits, bands appeared at expected molecular weights, with a notable exception being WaaA, which migrated approximately 15 kDa below its predicted size (47 kDa), likely due to anomalous SDS binding observed for some integral membrane proteins^23^. For the rest of hits, almost none had detectable bands with the ‘No SLB’ condition, while having measurable hits with at least one of the ‘DOPC SLB’ or ‘ECL SLB’ conditions (**Figure 2c**). The synthesis of other proteins, such as MscS, could only be observed with the addition of cardiolipin to the DOPC and ECL SLBs, which supports a previously characterized cardiolipin dependence of MscS^24^ (**Figure 2d**). The membrane-associated proteins SecA and PlsB, which demonstrated a preference for ECL SLBs in the initial screen, also interacted more strongly with a 1% addition of CL in the DOPC SLB, indicating a dependence on negatively charged lipids that can be resolved with ORB-TXTL.

Hit rates varied across functional classes, with notable enrichment among lipid synthesis enzymes, two-component histidine kinases, and ABC transporter ATPases (**Figure 2e**). However, the dominant predictor of hit rate was the number of transmembrane domains (TMDs): interacting membrane proteins (0 TMD), 1 TMD, and 2 TMD proteins showed hit rates of 83%, 67%, and 72%, respectively, while cytosolic proteins scored 10% (low as expected) and proteins with 5 or more TMDs scored 13% (**Figure 2f and 2g**). This sharp decline for proteins with 5 or more TMDs likely reflects one or more of the following: absence of membrane insertases in the cell-free system^25^, lack of a physiological transmembrane potential, or insufficient space on the outer leaflet closest to the silica surface for folding. To test the space hypothesis, PEGylated lipids were incorporated into the SLBs to create a spacer between the bilayer and the silica surface. For proteins with fewer TMDs, the interaction strengths decreased due to lower surface accessibility. Similarly, adding PEGylated lipids did not improve integration of multi-TMD proteins, with binding consistently being lower than the bare bead condition (**Figure S24**).

### ORB-TXTL can host a multi-enzyme bacterial lipid synthesis pathway

ORB-TXTL enables the functional reconstitution of multi-enzyme pathways by decoupling protein synthesis from catalytic activity: enzymes are first expressed together on SLB-coated beads, which are then washed and transferred to a reaction buffer for downstream assays (**Figure 3a**). Here, we applied this approach to the *E. coli* phospholipid synthesis pathway, beginning with determining the association to the bead’s SLBs of the pathway’s three enzymes – FadD, PlsB, and PlsC (**Figure 3b**). While FadD did not show a preference for either SLB composition, both PlsB and PlsC exhibited a mild preference for ECL SLBs. The *fadD* gene expressed multiple bands, suggesting alternative start codons (**Figure 3c**).

**Figure 3.**
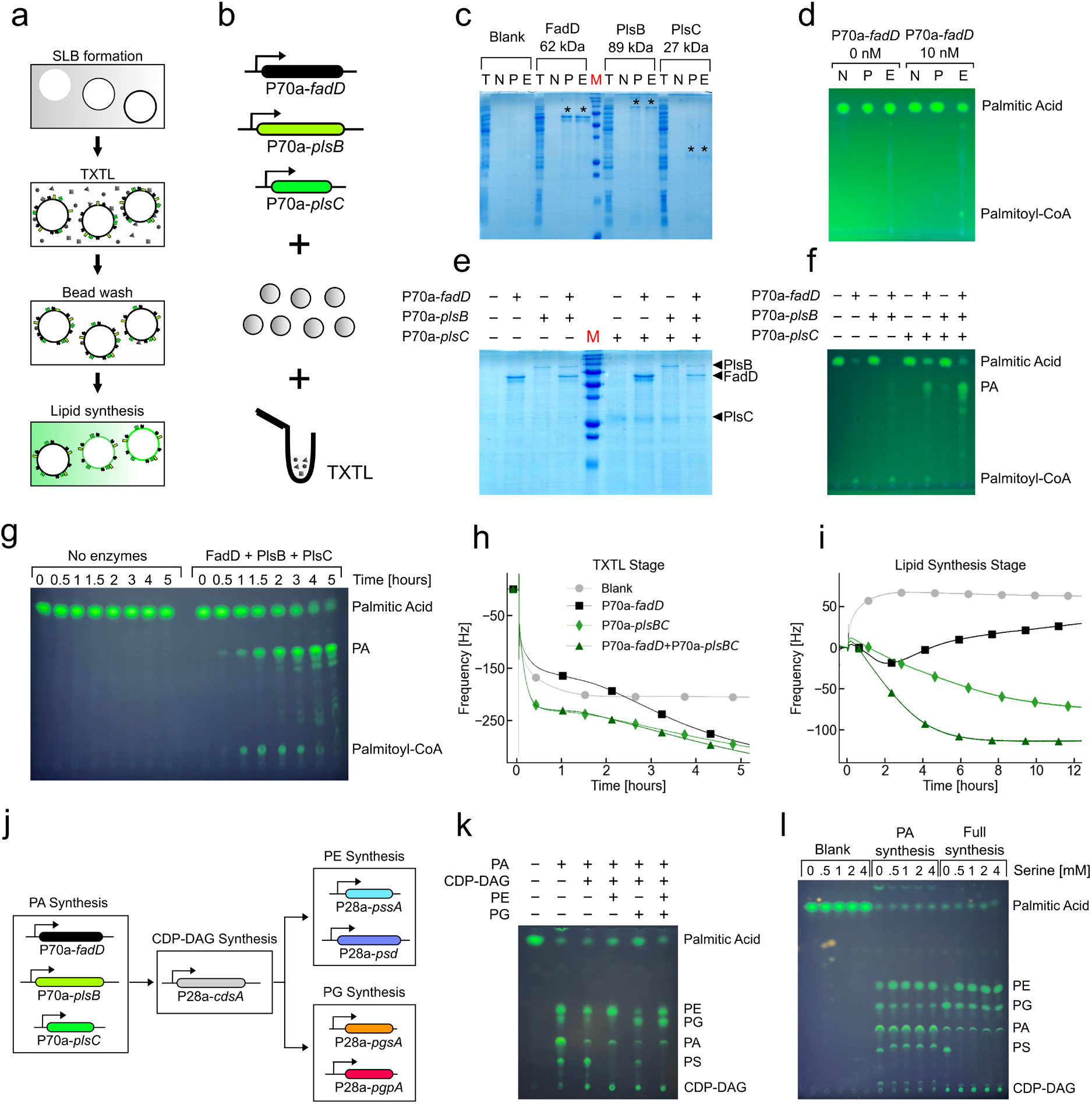
Two-stage cell-free biosynthesis of lipids on SLB-coated beads. **(a)** A schematic for the TXTL stage of the process, where some of the lipid synthesis membrane enzyme genes are expressed in contact with SLB-coated beads. **(b)** A schematic illustrating the two-stage workflow. **(c)** An SDS-PAGE gel showing the protein-lipid interactions of FadD, PlsB, and PlsC. **(d)** A TLC readout to track the synthesis of Palmitoyl-CoA, depending on whether FadD was synthesized and which SLB compositions were used. **(e)** An SDS-PAGE gel showing the interactions of FadD, PlsB, and PlsC with ECL SLB-coated beads when synthesized alone or in combinations. **(f)** A TLC readout to track the synthesis of Palmitoyl-CoA and PA for the various FadD, PlsB, and PlsC combinations on ECL SLB-coated beads. **(g)** A TLC readout for palmitoyl-CoA and PA synthesis over the duration of 5 hours when all enzymes are synthesized on ECL SLB-coated beads. **(h) and (i)** are the QCM-TXTL traces for the TXTL and Lipid Synthesis stages, respectively. **(j)** A schematic of the PE and PG lipid synthesis pathway broken up into genetic modules that can be expressed in TXTL. **(k)** A TLC readout demonstrating which lipids are synthesized depending on the combination of modules expressed in the TXTL stage. In the annotation above the TLC plate, each module is labeled by the lipid product. **(l)** A TLC readout comparing the synthesized lipids via the PA synthesis pathway or the full pathway with different Serine concentrations in the reaction.

Next, the activity of the phospholipid synthesis pathway’s enzymes was tested with fatty acid precursors. The primary substrate used was 90 µM palmitic acid, supplemented with 10 µM BODIPY-C16, a fluorescently labeled fatty acid^26^, to demonstrate phospholipid precursor specificity and increase assay sensitivity of incorporation of precursors into synthesized lipids. FadD activity was confirmed by TLC with both primuline and BODIPY detection. The conversion of fatty acids to acyl-CoA was only observed on ECL SLBs in the presence of TXTL-expressed *FadD*, which agrees with prior *in vitro* FadD studies (**Figure 3d, Figure S25**). Additionally, Acyl-CoA synthesis reached a steady state rather than accumulating, which was expected considering the reversibility of acyl-CoA thioesterification (**Figure S26**)^27^.

Afterwards, *fadD, plsB, and plsC* were expressed in all eight possible combinations on ECL SLB beads, and then the beads were washed and analyzed for the resulting ratios of synthesis concentrations (**Figure 3e**). Once their co-expression was confirmed, in another experiment, all three proteins were synthesized on ECL SLB beads, then the beads were washed and transferred to lipid synthesis buffer to determine phosphatidic acid synthesis. Phosphatidic acid was clearly detected by TLC as a BODIPY-tagged spot at the expected Rf value only when all three enzymes were present (**Figure 3f**). Since BODIPY-C16 is bulkier than palmitic acid, untagged fatty acids are expected to be consumed at equal or greater rates than BODIPY-C16, making the BODIPY signal a conservative estimate of flux. By this measure, the reaction reached approximately near complete substrate conversion after 5 hours (**Figure 3g**).

The phosphatidic acid synthesis assay TLC results were complemented by QCM measurements, which capture the kinetic signatures of lipid synthesis interacting with a sensor functionalized with an SLB with TXTL-expressed *fadD*, *plsB*, and *plsC*. Following protein synthesis, interfaced with an ECL SLB on a QCM, then washing (**Figure 3h**), lipid synthesis buffer 2 was introduced, and resonance frequency changes were monitored in real time (**Figure 3i**). The condition with all three enzymes showed a saturating frequency decrease, consistent with progressive lipid synthesis in the bilayer. FadD alone produced a transient frequency decrease followed by recovery, consistent with reversible acyl-CoA binding. The condition with PlsB/PlsC without FadD showed a gradual frequency decrease, which we attribute to acyl-CoAs present at low concentrations in the *E. coli* extract acting as substrates, adding perceived increases in mass on the sensor from associated acyl-CoA and converted phosphatidic acid.

Using the same approach as before, we functionalized the beads by synthesizing six different enzyme combinations that drive the pathway to produce: nothing, PA (PA synthesis module), CDP-DAG (PA and CDP-DAG synthesis modules), PE (PA, CDP-DAG, and PE synthesis modules), PG (PA, CDP-DAG, and PG synthesis modules), and PE with PG (PA, CDP-DAG, PE, and PG synthesis modules) (**Figure 3j**). In the PA-only condition, faint PE and PG bands were also visible, consistent with trace endogenous lipid synthesis enzymes present in the *E. coli* extract. Addition of CdsA dimmed the PA band and produced a CDP-DAG spot at Rf = 0.05, confirming flux through the pathway (**Figure 3k, Figure S27**). When the PE and PG modules were added, synthesis was funneled toward the respective products, though efficiency was lower when each module was expressed independently than when both were co-expressed, suggesting cooperative regulation between the two pathways. The synthesis of PE requires a displacement of the cytidine monophosphate (CMP) moiety with the amino acid serine, which produces the intermediate phospholipid PS. In the absence of serine in the lipid synthesis buffer, PE was not synthesized in both the ‘PA Synthesis’ and ‘Full Synthesis’ TXTL conditions (**Figure 3l, Figure S28**). However, increasing the serine concentration beyond 0.5 mM did not change the ratio of synthesized PE to PG, which again suggested regulation on the enzymatic level rather than substrate concentration^28^.

## Discussion

ORB-TXTL is a bead-based platform for rapid characterization of IIMPs lipid interactions and activity via CFE. Unlike liposomes or nanodiscs, beads can be extensively washed, transferred between reaction buffers, and read out by standard laboratory equipment – fluorescence microscopy, plate readers, SDS-PAGE, and TLC – without ultracentrifugation or specialized equipment. This makes ORB-TXTL accessible by laboratories working with IIMPs.

The principal limitation of ORB-TXTL is the synthesis of IIMPs with 4+ TMDs. The few polytopic hits observed in the screen all had relatively small and hydrophobic periplasmic loops, supporting the hypothesis that integration was spontaneous rather than active TMD shuttling^29^. Many valuable protein targets, such as GPCRs and transporters, have large critical extracytoplasmic domains that cannot fold without translocon assistance^30^. The functions of translocases and insertases were characterized *in vitro* via traditional purification and reconstitution approaches^31,32^, but the goal was mechanistic understanding. Nowadays, with advances in cell-free protein synthesis yields, decreasing DNA synthesis costs, and increasing demand for rapid membrane protein characterization, functional reconstitution of these pathways has again become a priority^33^. Future steps to address high-yield and general cell-free polytopic membrane protein synthesis should involve the integration and functionalization of beads with membrane protein insertases and translocon machinery.

Beyond targeted protein synthesis, we envision ORB-TXTL as a bottom-up systems biology platform. Here, we isolated single genes under a strong promoter-terminator pair, which is appropriate for systematic protein-lipid interaction screening. With improvements in large DNA sequencing and assembly^34,35^, similar experiments could be conducted with genomic sections on the order of 20-100 kb, enabling characterization of endogenous MP synthesis rates, their lipid interactions, and protein-protein interactions across entire membrane proteome segments. However, with parallel expression of many genes, detecting synthesis levels would require radioactive labeling^36^ and quantitative mass spectrometry readouts^37^.

## Methods

### Reagents

All the phospholipids were obtained from Avanti Polar Lipids in powder form. The phospholipids were dissolved in IPA (isopropyl alcohol, ThermoFisher Scientific, A416S-4) or 50:50 IPA:Chloroform mixture at the following stock concentrations: DOPC—20 mg/mL (840051 P), ECL (100500 P)—20 mg/mL, DOPE (1,2-dioleoyl-sn-glycero-3-phosphoethanolamine)—25 mg/mL in 50/50 chloroform/isopropanol (850725 P), DOPG (1,2-dioleoyl-sn-glycero-3-phospho-(1’-rac-glycerol))—50 mg/mL (840475 P), CL (1’,3’-bis[1,2-distearoyl-sn-glycero-3-phospho]-glycerol)—25 mg/mL (710334 P), DOPS (1,2-distearoyl-sn-glycero-3-phospho-L-serine) −12 mg/mL in 50/50 chloroform/isopropanol (840029 P), PEG5000-PE (1,2-dipalmitoyl-sn-glycero-3-phosphoethanolamine-N-[methoxy(polyethylene glycol)-5000])—0.5 mg/mL (880200 P). The fatty acid stocks were kept in ethanol: Oleic Acid – 1 M, Palmitic Acid – 20 mM, BODIPY-C16 (4,4-Difluoro-5,7-Dimethyl-4-Bora-3a,4a-Diaza-s-Indacene-3-Hexadecanoic Acid, D3821) – 2 mM. The cofactors used during lipid synthesis were all dissolved in water, flash frozen in liquid nitrogen, and stored at either −80°C (CoA) or at −20°C (G3P and L-Serine).

### Cell-free transcription-translation

CFE was carried out using an *E. coli* TXTL system described previously^25^, with one modification. We used the strain BL21-Δ*recBCD* Rosetta2 in which the *recBCD* gene set is knocked out to prevent the degradation of linear DNA. The preparation and usage of the TXTL system were the same as reported before. Briefly, *E. coli* cells were grown in a 2xYT medium supplemented with phosphates. Cells were pelleted, washed, and lysed with a cell press. After centrifugation, the supernatant was recovered and preincubated at 37 °C for 80 min. After a second centrifugation step, the supernatant was dialyzed for 3 h at 4 °C. After final centrifugation, the supernatant was aliquoted and stored at −80 °C. The TXTL reactions comprised the cell lysate, the energy and amino acid mixtures, maltodextrin (30 mM) and ribose (30 mM), magnesium (2–5 mM) and potassium (50–100 mM), PEG8000 (1–2 wt%), water, and the DNA to be expressed.

### Formation of supported lipid bilayers on silica beads

First, the bead powder (SBS Genetech, 5 µm Silica Microspheres) was mixed with water in a 10/90 volume ratio. If the stock was not used immediately, it was stored at 4°C for up to a month. For the washing stage, we dissolved 5 µL of the bead-water suspension in 1 mL of isopropanol in a 1.7 mL plastic tube. This suspension was vortexed and placed on a rotating platform (10 rpm) for 12 hours. After 12 hours, the tubes were agitated extensively by vortexing for 10 seconds, after which they were centrifuged for 10 seconds at 400 g with a benchtop mini centrifuge. The beads formed a pellet, and 950 µL of isopropanol was removed and replaced with 950 µL of fresh isopropanol (IPA). This IPA rinsing step was repeated four more times. After the final wash, the isopropanol volume was brought to an appropriate volume, and the lipids were added such that the beads were incubated in 1 mL of the lipid-IPA solution. For example, to incubate the beads in 3 mg/mL of ECL lipids, we added 150 µL of the 20 mg/mL ECL stock to 850 µL of the bead-isopropanol mixture. The beads vortexed to achieve a uniform lipid and bead distributions and were incubated for 25 minutes. Upon completion, the beads were spun down, IPA was removed such that the bead pellet remained submerged, and 900 µL of Saline Buffer 1 (pH 7.5, 150 mM NaCl, 10 mM Tris) was added. The beads were vortexed for 5 seconds at maximum speed to break down bead aggregation and then spun down again. The Saline Buffer was removed such that the beads remained submerged, and 900 µL of fresh buffer was added to the tube. This step was repeated two more times, such that the fraction of isopropanol in the sample was below 0.1%. This mixture was kept in the saline buffer until ready for the reaction.

### ORB-TXTL reactions

For most ORB-TXTL reactions described in this work, we used 15 µL of TXTL. First, the reactions were prepared as described above. Afterwards, the 1 mL beads mixture in the saline solution was vortexed, briefly spun down for 10s with a six-spot mini centrifuge, and then 980 µL of the buffer was removed, such that only a concentrated 20 µL suspension remained. To each 15 µL TXTL reaction, we added 2 µL of the suspension, which was consistently sufficient for 8 reactions. When the reactions were incubated in 1.7 mL tubes, we gently vortexed the solution until the beads were uniformly distributed and the reactions were placed in a thermoshaker programmed to shake for 12 hours at 24°C with a cycle of 1300 rpm for 1 minute and 700 rpm for 9 minutes. If the reactions were incubated in a 96-well plate, each condition was agitated via aspiration with a multichannel pipettor until the TXTL-bead suspension was uniform. The plate was sealed and taped to a platform rotating at 10 rpm for 12 hours. For experiments where the final readout was 10-well SDS PAGE, we used 25 µL of TXTL and 3 µL of the aforementioned concentrated 20 µL bead-buffer suspension. For lipid synthesis experiments, each condition used 90 µL of TXTL and 8 µL of concentrated bead-buffer suspension.

### Post-TXTL wash and sample preparation

After completion, each ORB-TXTL reaction is repeatedly treated with a wash buffer imitating the TXTL ionic composition (Hepes 50 mM, 10 mM Magnesium Glutamate, 150 mM Potassium Glutamate, pH 7.5). If the TXTL reaction was incubated in tubes, we added 400 µL of Wash Buffer, vortexed, spun down, removed 400 µL of the used wash buffer, and repeated two times. Afterwards, we add another 400 µL of the wash buffer and incubate the beads at room temperature on the bench. After 1 hour, we rinsed the buffer with 400 µL of the wash buffer two more times. The beads are then transferred to a fresh tube to prevent false signals from proteins attached to plastic walls. For MscL-mC fluorescence analysis, the beads were resuspended in 500 µL of the wash buffer, and 1 µL droplets were distributed on microscope slides, or 50 µL was dispensed in a 96-well plate. For SDS-PAGE experiments, all liquid wash was removed, and the beads were gently dried on the bench via evaporation. For the lipid synthesis reactions, the bead suspensions were concentrated to 10 µL in the reaction buffer. If the reactions were incubated in 96-well plates, to each well, we added 90 µL of the wash buffer. The plate was sealed and agitated by manual shaking. The plate was then spun down with a benchtop plate spinner, and the used wash buffer was removed. This step was repeated five times. After the 5^th^ rinse, we added 90 µL of fresh wash buffer and incubated the plate on a rotating platform for 1 hour. After the incubation, the beads were rinsed two more times, and the samples were transferred to a fresh 96-well plate. There, the beads were spun down, all supernatant was removed, and the wells were gently dried at room temperature on the bench.

### Fluorescence quantification with plate readers and microscopy, analysis

The Nile Red stains, Lact-C2 binding, and MscL integration experiments were imaged with an Olympus IX71 using 10x, 20x, and 40x lenses with the Texas Red Filter Cube (EX/EM 562/624 nm). For the Nile Red experiments, the exposure was set to 100 ms, while for the Lact-C2 and MscL experiments, the exposure was 250 ms. For analysis, the background was subtracted using a 100 px rolling ball method (beads were 13-15px). The beads were detected and traced using the ImageJ particle analysis software, and the traces were used as masks to collect fluorescent intensity data for each bead. Plate-based fluorescence measurements used Biotek H1 hybrid multi-mode plate readers. Experiments measuring mCherry and Rhodamine-PE fluorescence used 590 nm excitation 620 nm emission fluorescence.

### SDS-PAGE

Membrane protein interactions with SLB-coated beads were assessed by SDS-PAGE. Following the TXTL incubation and washing steps, beads were resuspended in SDS-PAGE sample buffer and incubated for 1 hour at room temperature to elute bound and integrated proteins. The eluate was loaded onto 12% or 15% polyacrylamide gels alongside a molecular weight ladder (Precision Plus Protein™ All Blue Prestained Protein Standards, BioRad #1610373). The electrophoresis was performed at 70 V for the stacking gel and 170 V for the resolving gel, which were cast in a Hoefer SE250 Mighty Small II Mini Vertical Protein Electrophoresis Unit. The gels were fixed in a methanol/acetic acid/water (40:10:50, v:v:v) solution for 10 hours, stained with 0.2% Coomassie in an ethanol/acetic acid/water (40:10:50, v:v:v) solution for 4 hours, and washed repeatedly in water for 24-72 hours. The gels were imaged with a document scanner (Epson Perfection V19 II).

### Lipid synthesis buffers

Each 100 µL ORB-TXTL reaction was washed as described above, concentrated to 10 µL, and added to 40 µL of the following lipid synthesis buffers in a fresh 1.7 mL tube. The reaction was vortexed and incubated in a thermoshaker at 37°C with a repeating cycle of 1 minute at 1300 rpm and 4 minutes at 700 rpm.

LSB-1: 50 mM Hepes, 7.5 pH, 10 mM Magnesium Glutamate, 50 mM Potassium Glutamate, 5 mM CoA, 1 mM Oleic Acid, 10 µM BODIPY-C16, 20-fold toolbox 2.0 diluted energy regeneration buffer^38^, water to 400 µL.

LSB-2: 50 mM Hepes, 7.5 pH, 10 mM Magnesium Glutamate, 50 mM Potassium Glutamate, 5 mM CoA, 90 µM Palmitic Acid, 10 µM BODIPY-C16, 20-fold toolbox 2.0 diluted energy regeneration buffer.

LSB-3: 50 mM Hepes, 7.5 pH, 10 mM Magnesium Glutamate, 50 mM Potassium Glutamate, 5 mM CoA, 90 µM Palmitic Acid, 10 µM BODIPY-C16, 20-fold toolbox 2.0 diluted energy regeneration buffer 1 mM G3P (for PA and PG synthesis).

LSB-4: 50 mM Hepes, 7.5 pH, 10 mM Magnesium Glutamate, 50 mM Potassium Glutamate, 5 mM CoA, 90 µM Palmitic Acid, 10 µM BODIPY-C16, 20-fold toolbox 2.0 diluted energy regeneration buffer, 1 mM G3P (for PA and PG synthesis), and 1 mM L-Serine (for PS and PE synthesis).

### Preparation of ORB-TXTL lipids synthesis samples for thin-layer chromatography

To the completed 50 µL reaction, we added 160 µL of methanol and 80 µL of chloroform. The sample was vortexed for 30 seconds, and 72 µL of 1% Glacial Acetic Acid solution was added. The sample was vortexed again and centrifuged at 5000 g for 3 minutes. The bottom phase containing the lipids was collected and placed in a 1 mL glass tube and sealed. The tubes were either analyzed immediately or stored at −20°C for up to two days.

### Thin Layer Chromatography

Lipid products were resolved by thin-layer chromatography on silica gel 60 plates. For all the experiments, the samples were fully dried under a nitrogen stream and resuspended with 10 µL of chloroform. For spotting, the samples were dispensed on the TLC plates in 2 µL at a time. For experiments resolving PA, CDP-DAG, PE, and PG, the samples were developed in chloroform/methanol/water (65:25:4, v/v/v). For panels resolving PA, Palmitic Acid, and Palmitoyl-CoA, the solvent system was adjusted to n-butanol/acetic acid/water (4:1:1, v/v/v). Lipid standards were run in parallel to assign Rf values. Plates were visualized by primuline staining under UV illumination or by direct fluorescence imaging of BODIPY-labeled species using a gel imager.

### QCM sensors and modules preparation

A QSense Analyzer (Biolin Scientific, Gothenburg, Sweden) with four channels was used with open modules to verify expression and association with SLBs formed on silicon dioxide sensors (Biolin Scientific, QSX303). The QCM measurements were done as previously described^18^, except with open modules. Sensors and modules were soaked in 1% SDS for 10 minutes, then soaked in ultrapure water for 10 minutes. Then, sensors and modules were rinsed with ultrapure water before being dried with nitrogen, and then the sensors were plasma-cleaned (Harrick Plasma, PDG-32) at low RF power under vacuum for 6 minutes. The sensors were then reinserted into the dry QCM modules. The QCM modules were closed and connected to the QSense Analyzer mount, and QSoft software was used to calibrate the sensors’ resonance frequencies and harmonics, with the sensors ready for experiments.

### QCM SLB preparation and TXTL reaction

The QCM modules were maintained at 22°C throughout the whole experiment. First, 930 µL of a Tris NaCl Buffer (10 mM Tris, 150 mM NaCl, pH 7.5) was added to all the modules until the resonance frequency of the sensors stabilized, which took 15-30 minutes. Next, the Tris Buffer was replaced with IPA via pipetting, and we waited for the signal to stabilize. Buffer or fluid replacement of each open module well in the QCM consisted of the following standard exchange (SE) protocol that left a minimum amount of liquid remaining above the sensor to avoid sensor dewetting: With liquid in the open module well, all liquid except 130 µL was removed and discarded, followed by an addition of 800 µL of the new liquid. This removal of 800 µL of old liquid and addition of 800 µL of new liquid was repeated three more times, such that 930 µL of new liquid remained after a total of four standard exchanges. The frequency shift for the Tris NaCl buffer to IPA SE step was −80 Hz. We then added lipids to the IPA and left them to incubate for 30 minutes for DOPC and 60 minutes for ECL. After stabilization, the lipid-IPA solution resonance frequency decreases by −5 to −10 Hz from the IPA solution. Next, we replaced the lipid-IPA solution via SE with the Tris NaCl buffer to complete the formation of the SLB. For DOPC SLBs, the frequency shift between a bare sensor and a sensor with an SLB is 25-28 Hz, while for ECL SLBs, the shift is 35-50 Hz. Finally, we replaced the Tris Buffer with the TXTL reactions, taped over the open module wells with transparent tape, and incubated the reactions in contact with the SLB-sensor system for 12-20 hours.

### QCM data analysis

The analysis of QSense Analyzer data was performed on the 7^th^ overtone of the resonance of the sensor due to its lowest sensitivity to variations in the mounting of the sensor, following the manufacturer’s recommendation. The frequency at the end of the second Tris NaCl flush during the SLB preparation was reset as Δf = 0. The frequency output of the seventh harmonic was divided by seven to reflect the effect the TXTL reactions had on the fundamental frequency changes of the sensor.

### Library construction with double PCR

First, the primers are selected for PCR extraction of the gene of interest. The 5’ primer starts with a CTTTAAGAAGGAGAGGTACCA overhang, while the 3’ primer starts with CGGCGGGCTTTGCTCGAG overhang. We perform the PCR reaction (NEB Q5) for 20-25 cycles with 0.2 µM of each primer and using frozen washed *E. coli* K12 cultures as the template. After the amplification was complete, we verified the amplicon length with DNA electrophoresis and PCR-cleaned the product. For the second round, we used the universal primers:

*P70a-UTR1-s2:* (Tm = 62) GTTCCGCTGGGCATGCTGAGCTAACACCGTGCGTGTTGACAATTTTACCTCTGGCGG TGATAATGGTTGCAGCTAGCAATAATTTTGTTTAACTTTAAGAAGGAGAGGTACCA ATG *T500-as1:* (Tm = 62.9)

GTCGACACAGAAAAGCCCGCCTTTCGGCGGGCTTTGCTCGAG

A 100 µL second round reaction (NEB Q5) used 25-50 ng of the washed first round PCR product as the template and 0.35 µM of the universal primers. The annealing and extension temperatures for the first 10 cycles were 68°C and 72°C, whilst for the last 25 cycles, the temperatures of both were raised to 76°C. The increased temperature reduced the efficiency of the polymerase, thus the extension times were raised to 45 seconds per 1 kbp, instead of the recommended 20-30 seconds. Upon completion, the construct length was validated with DNA electrophoresis, and amplified DNA was PCR purified using a homemade kit. All DNA sequences used in this work are organized in the file titled ‘2026_ORB_DNA’.

## Data Availability

The data used in Figures 1-3 and Supplementary Figures 1-28 are provided as source data files in Supplementary Data 1. Plasmids and linear DNA used in this study are listed in Supplementary Data 2.

## Author Contributions

A.K. and S.T. performed the experiments and analyzed the data. A.K. developed the methodology. A.K., S.T., and V.N. designed the experiments and wrote the manuscript. V.N. edited the manuscript, supervised the project, and acquired funding.

## Acknowledgements

The authors thank David Garenne and Paul Soudier for their help in the preparation of the TXTL system used in this work. This work and the materials are based on funding provided by the National Science Foundation (BBSRC-NSF/BIO 2017932 to V.N.).

## References

(1) Alberts, B.; Johnson, A.; Lewis, J.; Raff, M.; Roberts, K.; Walter, P. Membrane Proteins. In Molecular Biology of the Cell. 4th edition; Garland Science, 2002.

(2) Schneider, B.; Junge, F.; Shirokov, V. A.; Durst, F.; Schwarz, D.; Dötsch, V.; Bernhard, F. Membrane Protein Expression in Cell-Free Systems. In Heterologous Expression of Membrane Proteins: Methods and Protocols; Mus-Veteau, I., Ed.; Humana Press: Totowa, NJ, 2010; pp 165–186. 10.1007/978-1-60761-344-2_11.

(3) Kuruma, Y.; Ueda, T. The PURE System for the Cell-Free Synthesis of Membrane Proteins. Nat Protoc 2015, 10 (9), 1328–1344. 10.1038/nprot.2015.082.

(4) Membrane protein synthesis: no cells required: Trends in Biochemical Sciences. https://www.cell.com/trends/biochemical-sciences/abstract/S0968-0004(23)00082-8 (accessed 2026-06-24).

(5) Niwa, T.; Sasaki, Y.; Uemura, E.; Nakamura, S.; Akiyama, M.; Ando, M.; Sawada, S.; Mukai, S.; Ueda, T.; Taguchi, H.; Akiyoshi, K. Comprehensive Study of Liposome-Assisted Synthesis of Membrane Proteins Using a Reconstituted Cell-Free Translation System. Sci Rep 2015, 5 (1), 18025. 10.1038/srep18025.

(6) Meyer, C.; Arizzi, A.; Henson, T.; Aviran, S.; Longo, M. L.; Wang, A.; Tan, C. Designer Artificial Environments for Membrane Protein Synthesis. Nat Commun 2025, *16* (1), 4363. 10.1038/s41467-025-59471-1.

(7) Henson, T.; Arizzi, A.; Meyer, C.; Wang, D.; Lowe, N. M.; Wang, Y.; Ananda, K.; Carney, R. P.; Wang, A.; Tan, C. Prototyping Minimal Extracellular Vesicle Mimetics Using Cell-Free Synthesis. ACS Nano 2026, 20 (9), 7385–7400. 10.1021/acsnano.5c05047.

(8) Coleman, M. A.; Cappuccio, J. A.; Blanchette, C. D.; Gao, T.; Arroyo, E. S.; Hinz, A. K.; Bourguet, F. A.; Segelke, B.; Hoeprich, P. D.; Huser, T.; Laurence, T. A.; Motin, V. L.; Chromy, B. A. Expression and Association of the Yersinia Pestis Translocon Proteins, YopB and YopD, Are Facilitated by Nanolipoprotein Particles. PLOS ONE 2016, 11 (3), e0150166. 10.1371/journal.pone.0150166.

(9) Bruni, R.; Laguerre, A.; Kaminska, A.-M.; McSweeney, S.; Hendrickson, W. A.; Liu, Q. High-Throughput Cell-Free Screening of Eukaryotic Membrane Protein Expression in Lipidic Mimetics. Protein Science 2022, 31 (3), 639–651. 10.1002/pro.4259.

(10) Köck, Z.; Schnelle, K.; Persechino, M.; Umbach, S.; Schihada, H.; Januliene, D.; Parey, K.; Pockes, S.; Kolb, P.; Dötsch, V.; Möller, A.; Hilger, D.; Bernhard, F. Cryo-EM Structure of Cell-Free Synthesized Human Histamine 2 Receptor/Gs Complex in Nanodisc Environment. Nat Commun 2024, 15 (1), 1831. 10.1038/s41467-024-46096-z.

(11) Peruzzi, J. A.; Vu, T. Q.; Gunnels, T. F.; Kamat, N. P. Rapid Generation of Therapeutic Nanoparticles Using Cell-Free Expression Systems. Small Methods 2023, 7 (12), 2201718. 10.1002/smtd.202201718.

(12) Klammt, C.; Schwarz, D.; Eifler, N.; Engel, A.; Piehler, J.; Haase, W.; Hahn, S.; Dötsch, V.; Bernhard, F. Cell-Free Production of G Protein-Coupled Receptors for Functional and Structural Studies. Journal of Structural Biology 2007, 158 (3), 482–493. 10.1016/j.jsb.2007.01.006.

(13) Ma, Y.; Münch, D.; Schneider, T.; Sahl, H.-G.; Bouhss, A.; Ghoshdastider, U.; Wang, J.; Dötsch, V.; Wang, X.; Bernhard, F. Preparative Scale Cell-Free Production and Quality Optimization of MraY Homologues in Different Expression Modes *. Journal of Biological Chemistry 2011, *286* (45), 38844–38853. 10.1074/jbc.M111.301085.

(14) Focke, P. J.; Hein, C.; Hoffmann, B.; Matulef, K.; Bernhard, F.; Dötsch, V.; Valiyaveetil, F. I. Combining in Vitro Folding with Cell Free Protein Synthesis for Membrane Protein Expression. Biochemistry 2016, 55 (30), 4212–4219. 10.1021/acs.biochem.6b00488.

(15) Lu, Y.; Allegri, G.; Huskens, J. Vesicle-Based Artificial Cells: Materials, Construction Methods and Applications. Mater. Horiz. 2022, 9 (3), 892–907. 10.1039/d1mh01431e.

(16) Neumann, S.; Pucadyil, T. J.; Schmid, S. L. Analyzing Membrane Remodeling and Fission Using Supported Bilayers with Excess Membrane Reservoir. Nat Protoc 2013, 8 (1), 213–222. 10.1038/nprot.2012.152.

(17) Majumder, S.; Willey, P. T.; DeNies, M. S.; Liu, A. P.; Luxton, G. W. G. A Synthetic Biology Platform for the Reconstitution and Mechanistic Dissection of LINC Complex Assembly. J Cell Sci 2018, 132 (4), jcs219451. 10.1242/jcs.219451.

(18) Khakimzhan, A.; Izri, Z.; Thompson, S.; Dmytrenko, O.; Fischer, P.; Beisel, C.; Noireaux, V. Cell-Free Expression with a Quartz Crystal Microbalance Enables Rapid, Dynamic, and Label-Free Characterization of Membrane-Interacting Proteins. Commun Biol 2024, 7 (1), 1005. 10.1038/s42003-024-06690-9.

(19) Ferhan, A. R.; Yoon, B. K.; Park, S.; Sut, T. N.; Chin, H.; Park, J. H.; Jackman, J. A.; Cho, N.-J. Solvent-Assisted Preparation of Supported Lipid Bilayers. Nat Protoc 2019, 14 (7), 2091–2118. 10.1038/s41596-019-0174-2.

(20) Tabaei, S. R.; Vafaei, S.; Cho, N.-J. Fabrication of Charged Membranes by the Solvent-Assisted Lipid Bilayer (SALB) Formation Method on SiO2 and Al2O3. Phys. Chem. Chem. Phys. 2015, 17 (17), 11546–11552. 10.1039/C5CP01428J.

(21) Andersen, M. H.; Graversen, H.; Fedosov, S. N.; Petersen, T. E.; Rasmussen, J. T. Functional Analyses of Two Cellular Binding Domains of Bovine Lactadherin. Biochemistry 2000, 39 (20), 6200–6206. 10.1021/bi992221r.

(22) Haswell, E. S.; Phillips, R.; Rees, D. C. Mechanosensitive Channels: What Can They Do and How Do They Do It? Structure 2011, 19 (10), 1356–1369. 10.1016/j.str.2011.09.005.

(23) Rath, A.; Glibowicka, M.; Nadeau, V. G.; Chen, G.; Deber, C. M. Detergent Binding Explains Anomalous SDS-PAGE Migration of Membrane Proteins. Proceedings of the National Academy of Sciences 2009, 106 (6), 1760–1765. 10.1073/pnas.0813167106.

(24) Ridone, P.; Nakayama, Y.; Martinac, B.; Battle, A. R. Patch Clamp Characterization of the Effect of Cardiolipin on MscS of E. Coli. Eur Biophys J 2015, 44 (7), 567–576. 10.1007/s00249-015-1020-2.

(25) Garenne, D.; Thompson, S.; Brisson, A.; Khakimzhan, A.; Noireaux, V. The All-E. Coli TXTL Toolbox 3.0: New Capabilities of a Cell-Free Synthetic Biology Platform. Synth Biol 2021, 6 (1), ysab017. 10.1093/synbio/ysab017.

(26) Loudet, A.; Burgess, K. BODIPY Dyes and Their Derivatives: Syntheses and Spectroscopic Properties. Chem. Rev. 2007, 107 (11), 4891–4932. 10.1021/cr078381n.

(27) FÆrgeman, N. J.; Knudsen, J. Role of Long-Chain Fatty Acyl-CoA Esters in the Regulation of Metabolism and in Cell Signalling. Biochem J 1997, 323 (1), 1–12. 10.1042/bj3230001.

(28) Lee, E.; Cho, G.; Kim, J. Structural Basis for Membrane Association and Catalysis by Phosphatidylserine Synthase in Escherichia Coli. Science Advances 2024, 10 (51), eadq4624. 10.1126/sciadv.adq4624.

(29) Engelman, D. M.; Steitz, T. A. The Spontaneous Insertion of Proteins into and across Membranes: The Helical Hairpin Hypothesis. Cell 1981, 23 (2), 411–422. 10.1016/0092-8674(81)90136-7.

(30) Hegde, R. S.; Keenan, R. J. A Unifying Model for Membrane Protein Biogenesis. Nat Struct Mol Biol 2024, 31 (7), 1009–1017. 10.1038/s41594-024-01296-5.

(31) Brundage, L.; Hendrick, J. P.; Schiebel, E.; Driessen, A. J. M.; Wickner, W. The Purified E. Coli Integral Membrane Protein SecY E Is Sufficient for Reconstitution of SecA-Dependent Precursor Protein Translocation. Cell 1990, 62 (4), 649–657. 10.1016/0092-8674(90)90111-Q.

(32) Samuelson, J. C.; Chen, M.; Jiang, F.; Möller, I.; Wiedmann, M.; Kuhn, A.; Phillips, G. J.; Dalbey, R. E. YidC Mediates Membrane Protein Insertion in Bacteria. Nature 2000, 406 (6796), 637–641. 10.1038/35020586.

(33) Meier, M.; Scholz, S. A.; Bank, L. von; Levandoski, J. E.; Lückhof, M.; Schaaf, M.; Si, S.; Lieberwirth, I.; Jung, A. L.; Landfester, K.; Driessen, A. J. M.; Erb, T. J. Functional Mapping and Engineering of the Sec Translocon Unlocked by a Cell-Free System. bioRxiv December 11, 2025, p 2025.12.09.688994. 10.64898/2025.12.09.688994.

(34) Moss, E. L.; Maghini, D. G.; Bhatt, A. S. Complete, Closed Bacterial Genomes from Microbiomes Using Nanopore Sequencing. Nat Biotechnol 2020, 38 (6), 701–707. 10.1038/s41587-020-0422-6.

(35) Blount, B. A.; Lu, X.; Driessen, M. R. M.; Jovicevic, D.; Sanchez, M. I.; Ciurkot, K.; Zhao, Y.; Lauer, S.; McKiernan, R. M.; Gowers, G.-O. F.; Sweeney, F.; Fanfani, V.; Lobzaev, E.; Palacios-Flores, K.; Walker, R. S. K.; Hesketh, A.; Cai, J.; Oliver, S. G.; Cai, Y.; Stracquadanio, G.; Mitchell, L. A.; Bader, J. S.; Boeke, J. D.; Ellis, T. Synthetic Yeast Chromosome XI Design Provides a Testbed for the Study of Extrachromosomal Circular DNA Dynamics. Cell Genomics 2023, 3 (11). 10.1016/j.xgen.2023.100418.

(36) Kigawa, T.; Yabuki, T.; Yoshida, Y.; Tsutsui, M.; Ito, Y.; Shibata, T.; Yokoyama, S. Cell-Free Production and Stable-Isotope Labeling of Milligram Quantities of Proteins. FEBS Letters 1999, 442 (1), 15–19. 10.1016/S0014-5793(98)01620-2.

(37) Kirkpatrick, D. S.; Gerber, S. A.; Gygi, S. P. The Absolute Quantification Strategy: A General Procedure for the Quantification of Proteins and Post-Translational Modifications. Methods 2005, 35 (3), 265–273. 10.1016/j.ymeth.2004.08.018.

(38) Garamella, J.; Marshall, R.; Rustad, M.; Noireaux, V. The All E. Coli TX-TL Toolbox 2.0: A Platform for Cell-Free Synthetic Biology. ACS Synth. Biol. 2016, 5 (4), 344–355. 10.1021/acssynbio.5b00296.

