## supplementary information for "ORB-TXTL: cell-free expression of membrane proteins on lipid bilayer-coated beads"

Supplementary Figure 1: Composite Nile Red images of DOPC lipid ranges.
Supplementary Figure 2: Composite Nile Red images of ECL lipid ranges.
Supplementary Figure 3: Effect of lipid concentration on SLB density on beads and with the QCMD.
Supplementary Figure 4. Achieving SLB coverage for minimal non-specific TXTL adsorption on QCM sensors.
Supplementary Figure 5: Fluorescent images of Rhod-PE addition to DOPC SLBs.
Supplementary Figure 6: Improving MscL integration and fluorescent signal by diluting fluorescently tagged mutants.
Supplementary Figure 7: Composite images of MscL integration onto Bare, DOPC, and ECL coated beads.
Supplementary Figure 8: Detailed schematic of double PCR amplification.
Supplementary Figure 9: SDS-PAGE Gels Group 1.
Supplementary Figure 10: SDS-PAGE Gels Group 2.
Supplementary Figure 11: SDS-PAGE Gels Group 3.
Supplementary Figure 12: SDS-PAGE Gels Group 4.
Supplementary Figure 13: SDS-PAGE Gels Group 5.
Supplementary Figure 14: SDS-PAGE Gels Group 6.
Supplementary Figure 15: SDS-PAGE Gels Group 7.
Supplementary Figure 16: SDS-PAGE Gels Group 8.
Supplementary Figure 17: SDS-PAGE Gels Group 9.
Supplementary Figure 18: SDS-PAGE Gels Group 10.
Supplementary Figure 19: SDS-PAGE Gels Group 11.
Supplementary Figure 20: Gel Group 12. SDS-PAGE for MscS, CorA, and Kch.
Supplementary Figure 21: SDS-PAGE for combinations of P70a-*secY,* P70a-*yidC*, and P70a-*secA*.
Supplementary Figure 22: SDS-PAGE for combinations of P70a-*ftsY* and P70a-*ffh*.
Supplementary Figure 23: Effect of CL on the binding for a selection of proteins.
Supplementary Figure 24: Effect of PEG2k-PE on the binding for a selection of proteins. PEG2k-PE is added to ECL SLBs.
Supplementary Figure 25: Visualization of FadD activity with Primulin staining.
Supplementary Figure 26: Time course of palmitoyl-CoA synthesis.
Supplementary Figure 27: Visualization of lipid synthesis combinations with Primulin staining.
Supplementary Figure 28: L-Serine range for partial and full lipid synthesis pathways.

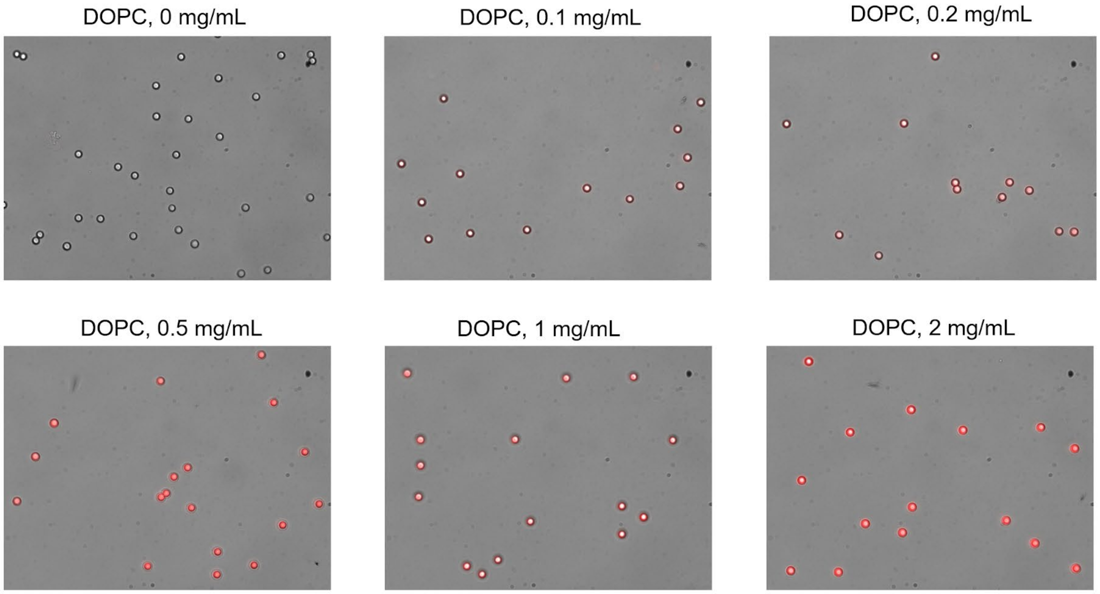


**Supplementary Figure 1.** Composite Nile Red images (phase contrast and fluorescence red channel) of DOPC lipid ranges on 5-micron beads. After forming the SLBs, the beads were incubated with 20 μM of Nile Red for 1 hour.

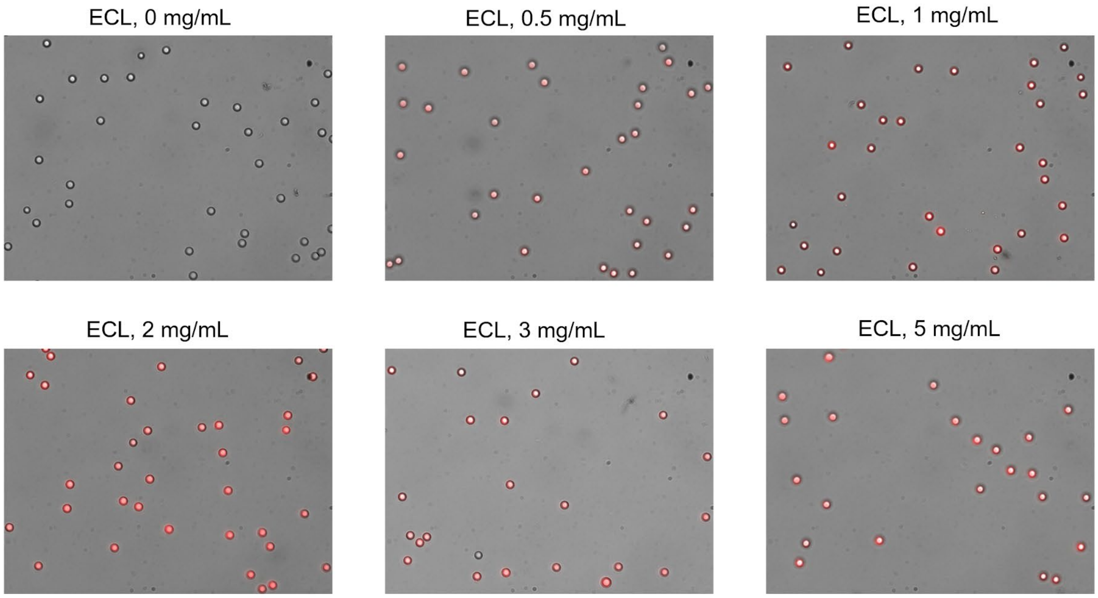


**Supplementary Figure 2.** Composite Nile Red images (phase contrast and fluorescence red channel) of ECL lipid ranges on 5-micron beads. After forming the SLBs, the beads were incubated with 20 μM of Nile Red for 1 hour.

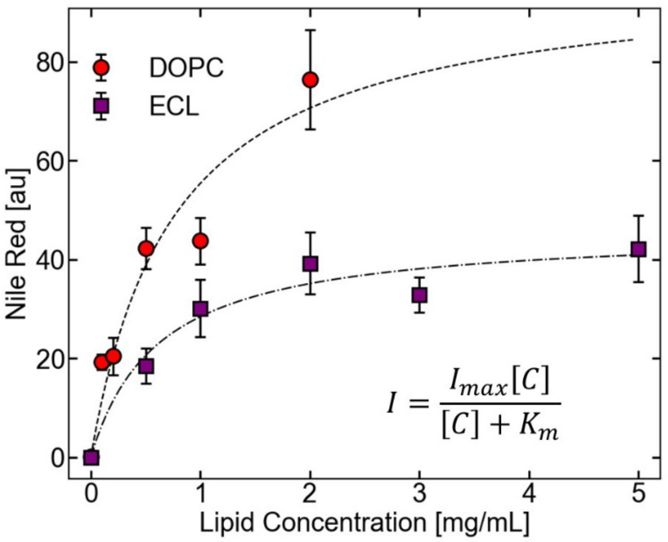


**Supplementary Figure 3.** Effect of lipid concentration on SLB density. The data were fit to a Hill function.

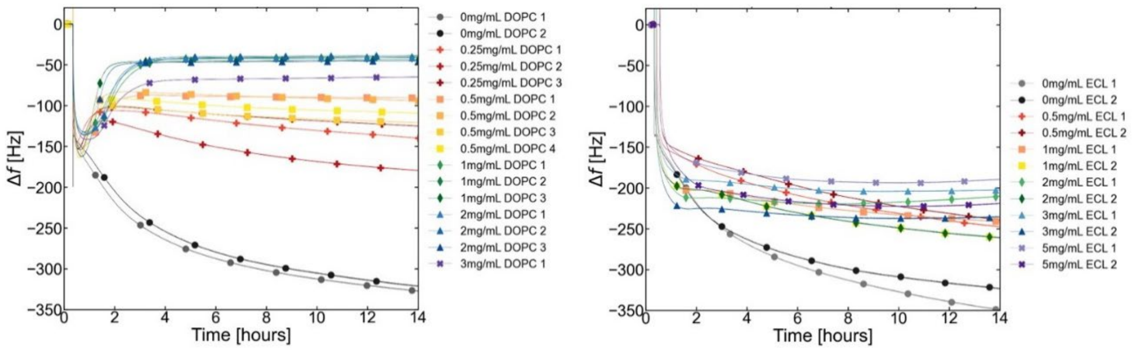


**Supplementary Figure 4.** Achieving SLB coverage for minimal non-specific TXTL adsorption on QCM sensors. Non-specific adsorption profiles of TXTL to the QCM sensor-SLB systems as a function of DOPC (right) and ECL (left) lipid concentrations in during SALB.

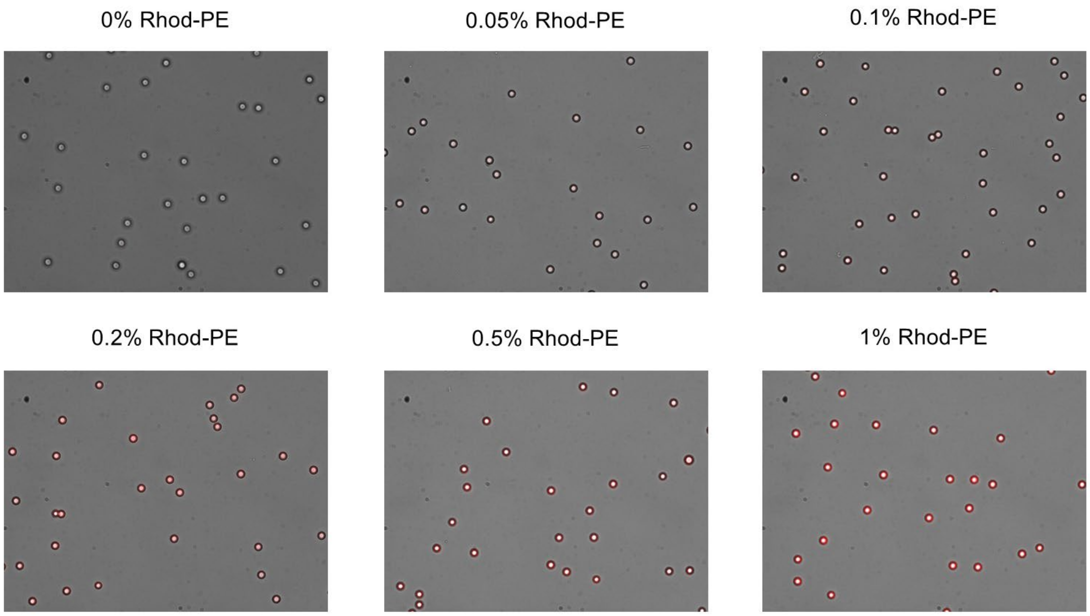


**Supplementary Figure 5.** Composite images of Rhod-PE added to DOPC SLBs on 5-micron beads.

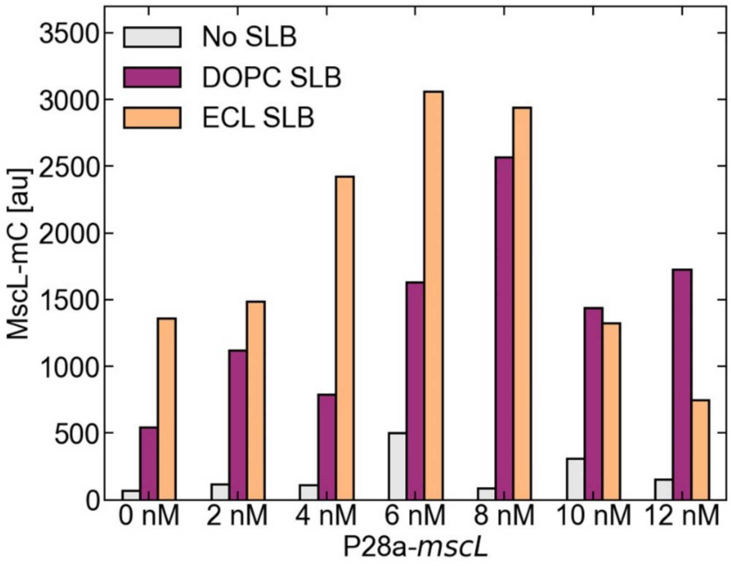


**Supplementary Figure 6.** Beads with No, DOPC, or ECL SLBs were added to TXTL reactions expressing 0.5 nM P70a-*s28* and 1 nM P28a-*mscL-mC*, over a range of P28a-*mscL* concentrations. This was to determine the optimum MscL integration and fluorescence signal by diluting fluorescently tagged mutants. Fluorescence measurements were obtained in a plate reader measuring mCherry signal (Ex/Em 560/610).

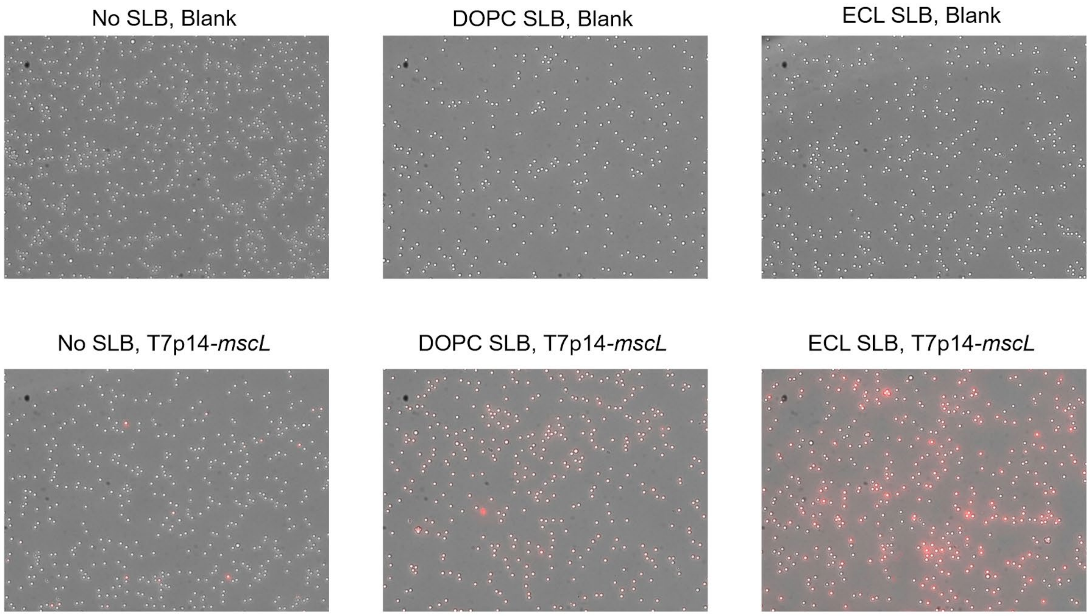


**Supplementary Figure 7.** Composite images (phase contrast and fluorescence red channel) of MscL integration onto Bare, DOPC, and ECL-coated 5-micron beads.

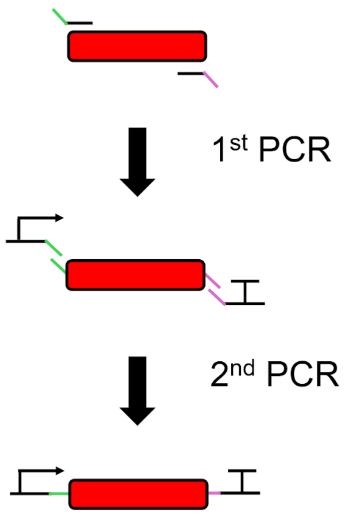


**Supplementary Figure 8.** Gene extraction protocol. First, the gene is extracted with overhang primers. After purification of the 1st PCR, the extracted DNA is used as a template for a second round, using 'universal primers' that contain a promoter and a terminator in the sense and antisense directions, respectively.

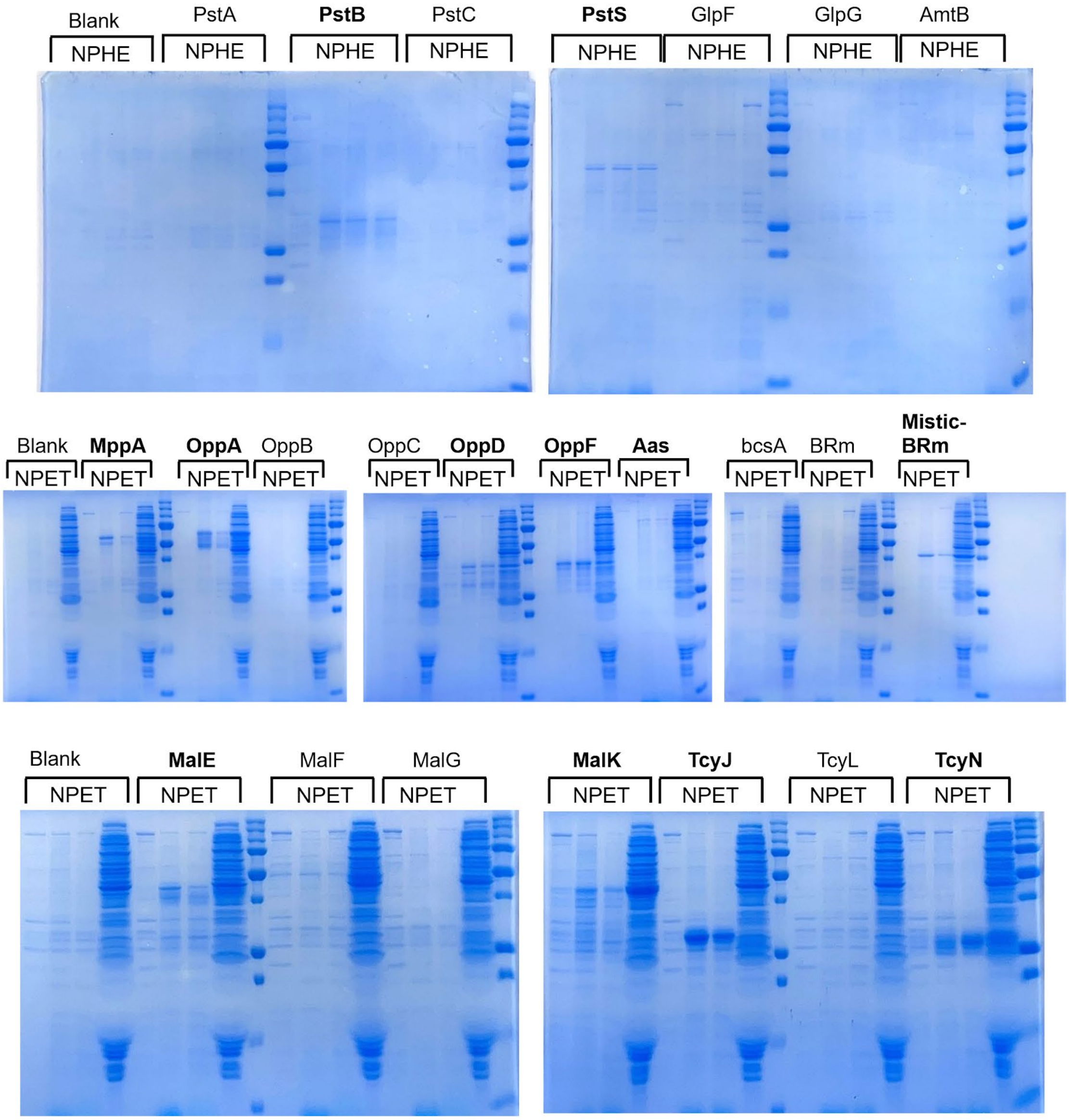


**Supplementary Figure 9.** SDS-PAGE Gels Group 1.

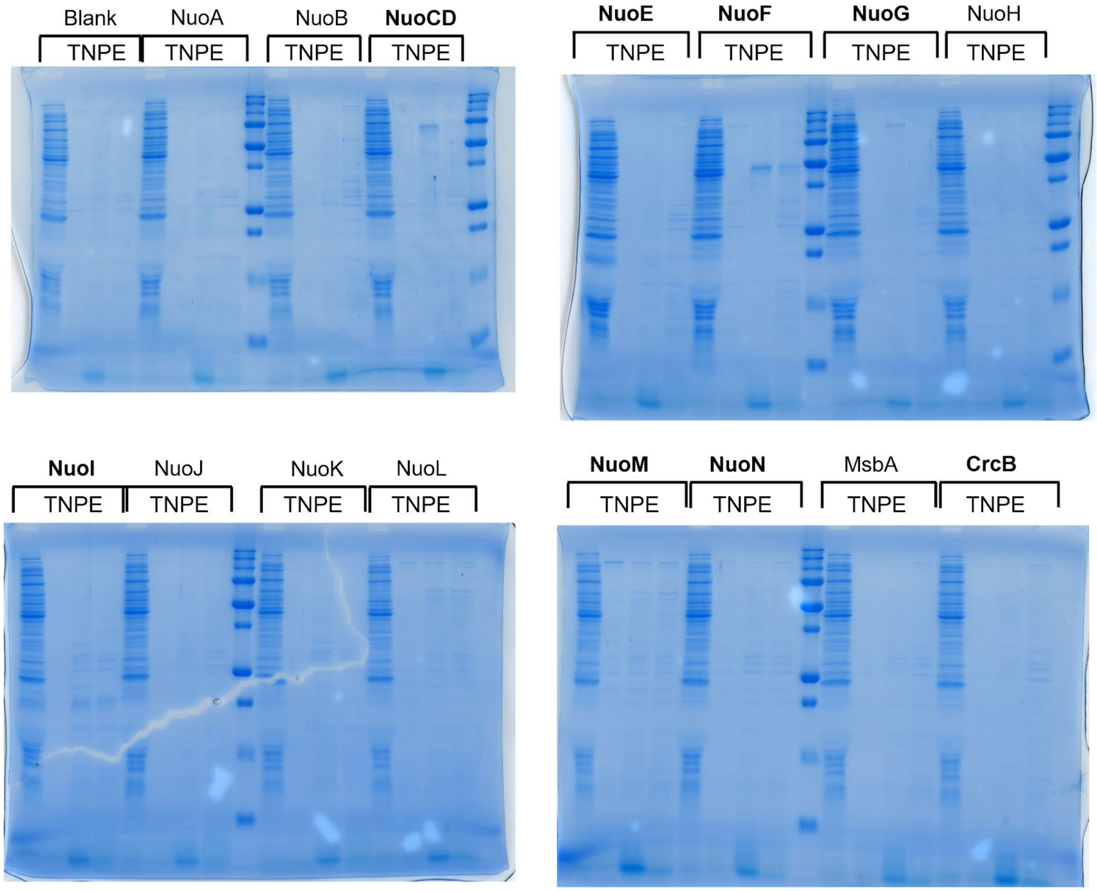


**Supplementary Figure 10.** SDS-PAGE Gels Group 2.

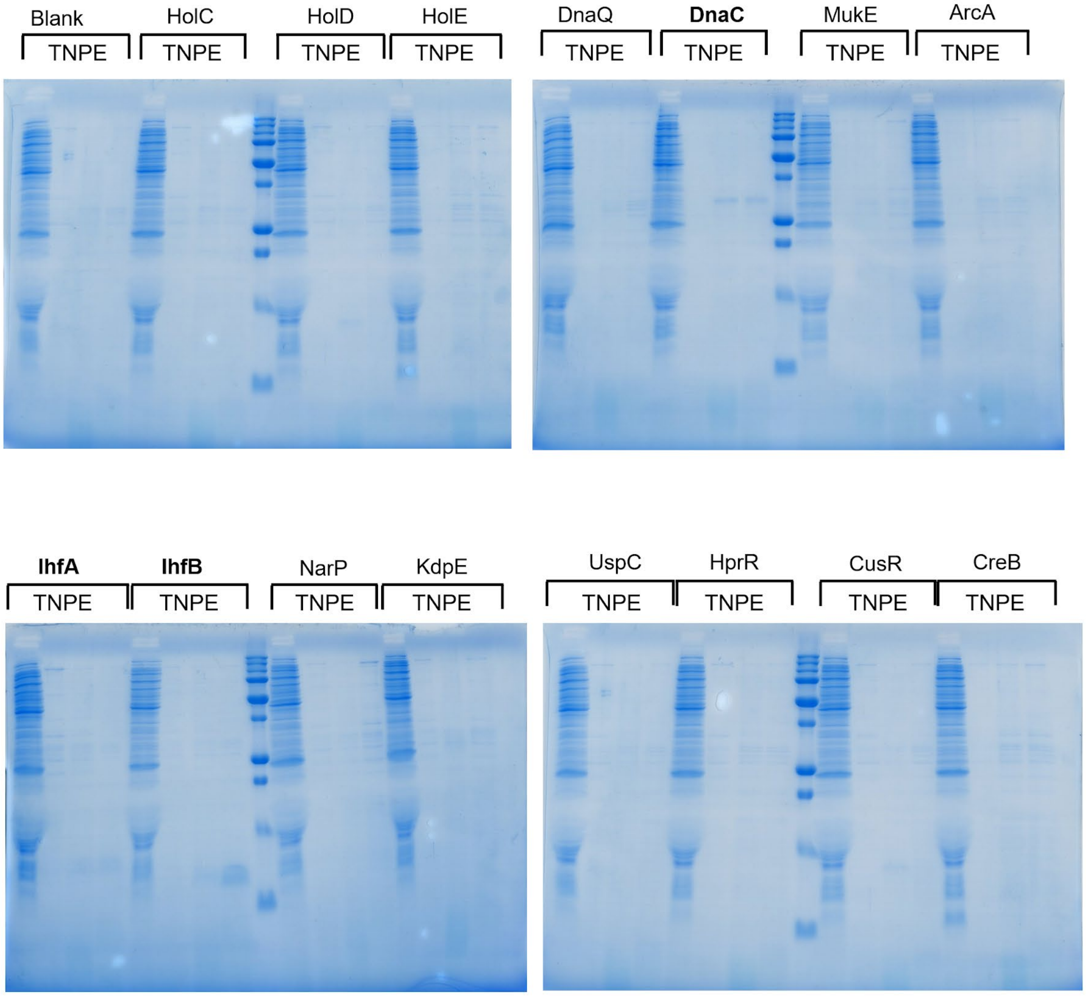


**Supplementary Figure 11.** SDS-PAGE Gels Group 3.

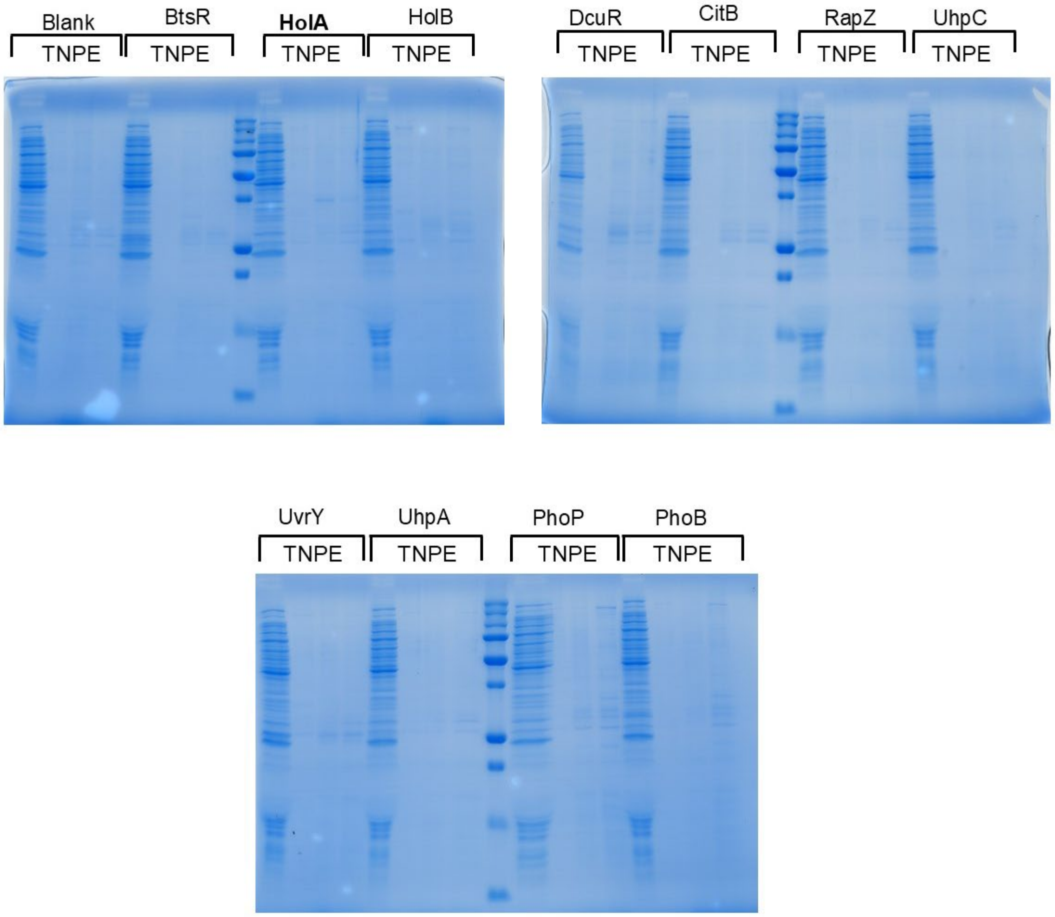


**Supplementary Figure 12.** SDS-PAGE Gels Group 4.

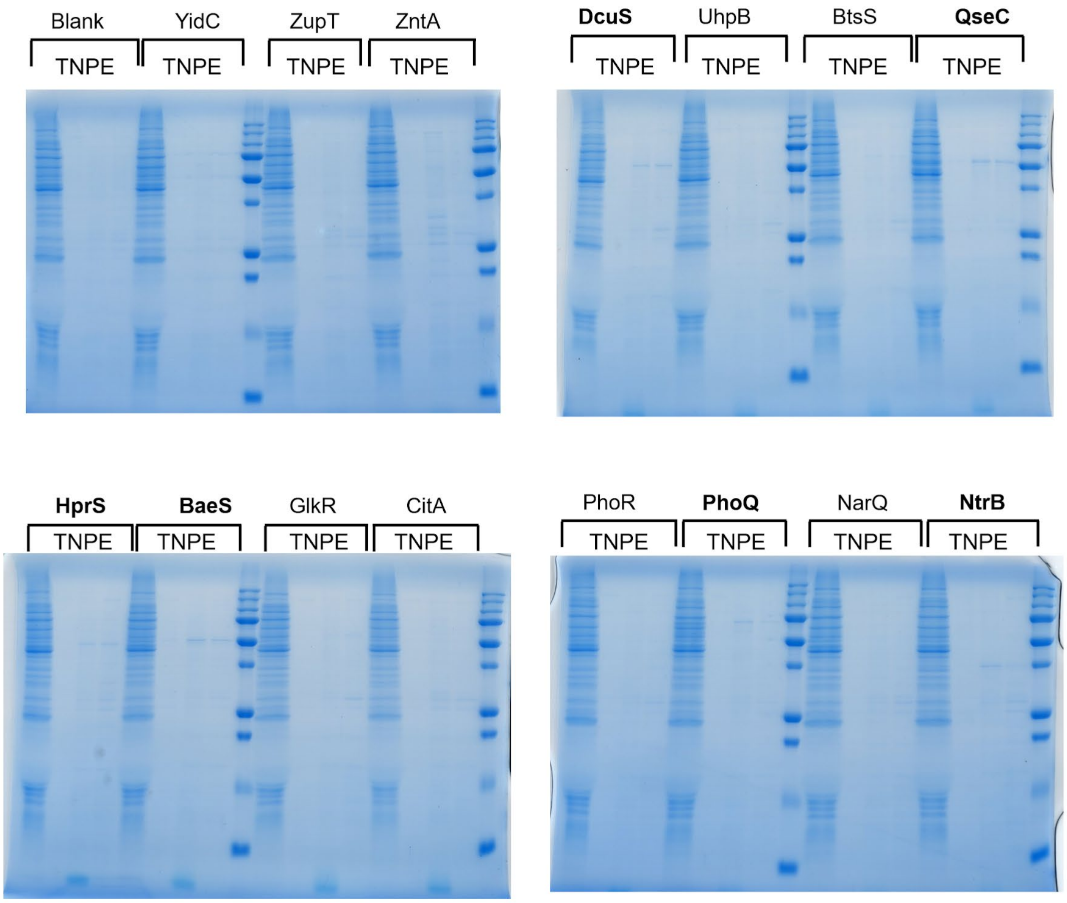


**Supplementary Figure 13.** SDS-PAGE Gels Group 5.

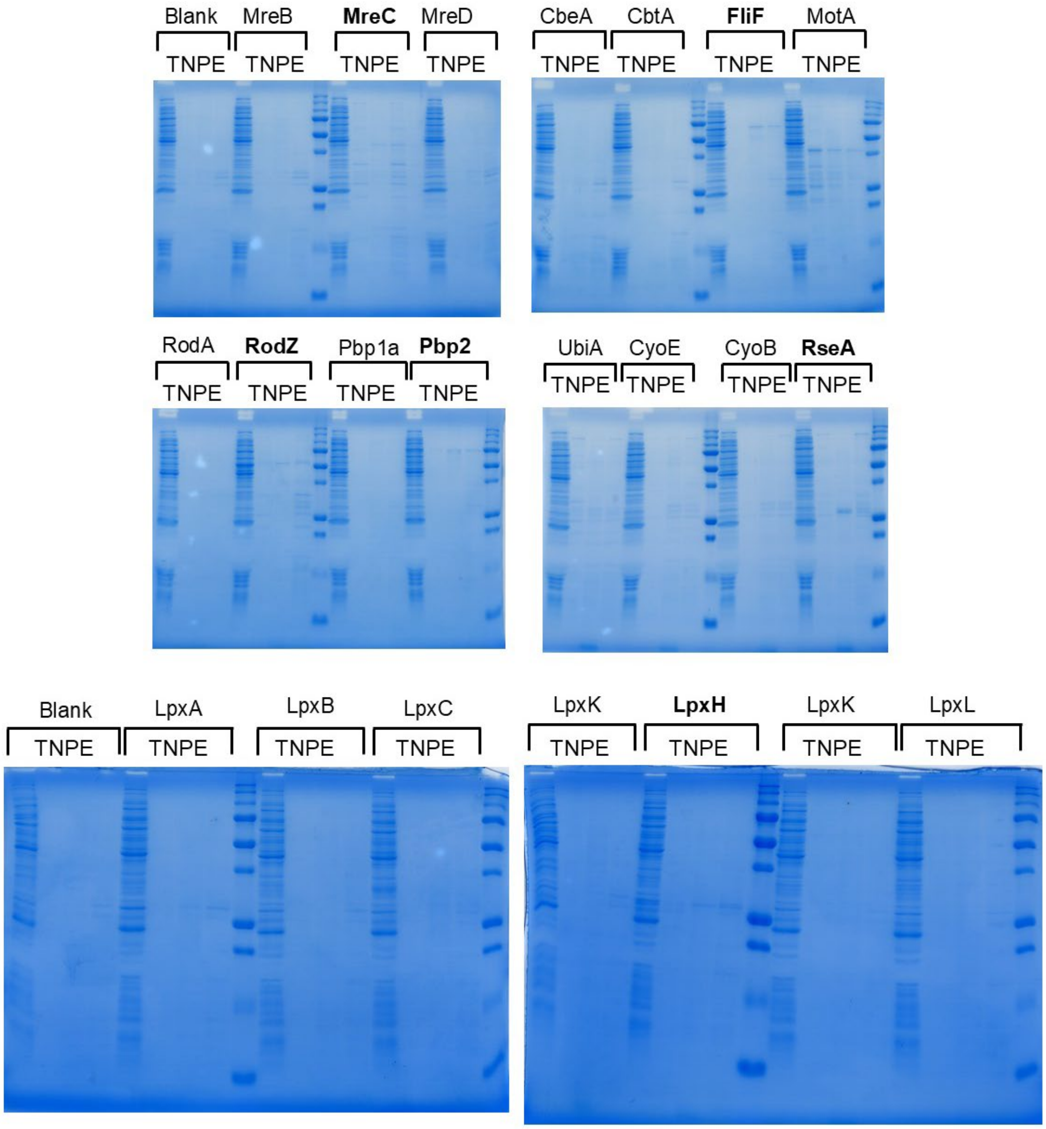


**Supplementary Figure 14.** SDS-PAGE Gels Group 6.

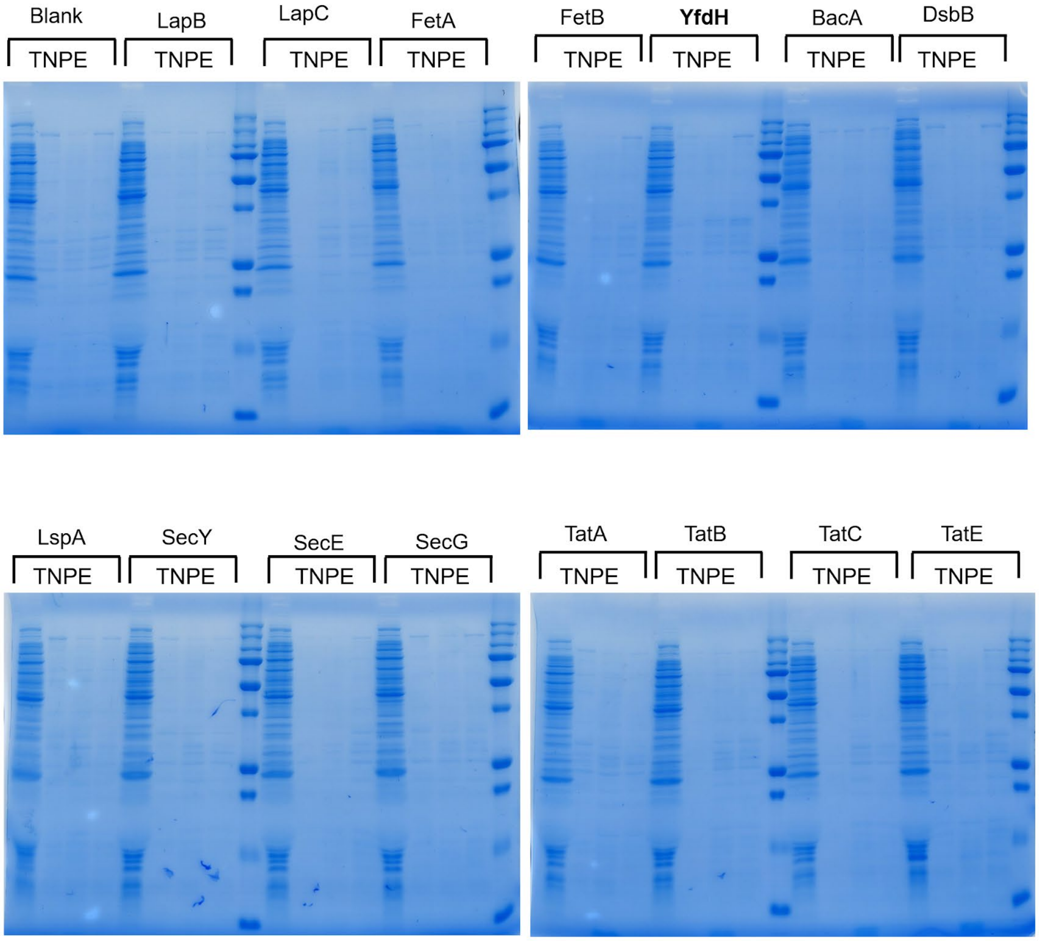


**Supplementary Figure 15.** SDS-PAGE Gels Group 7.

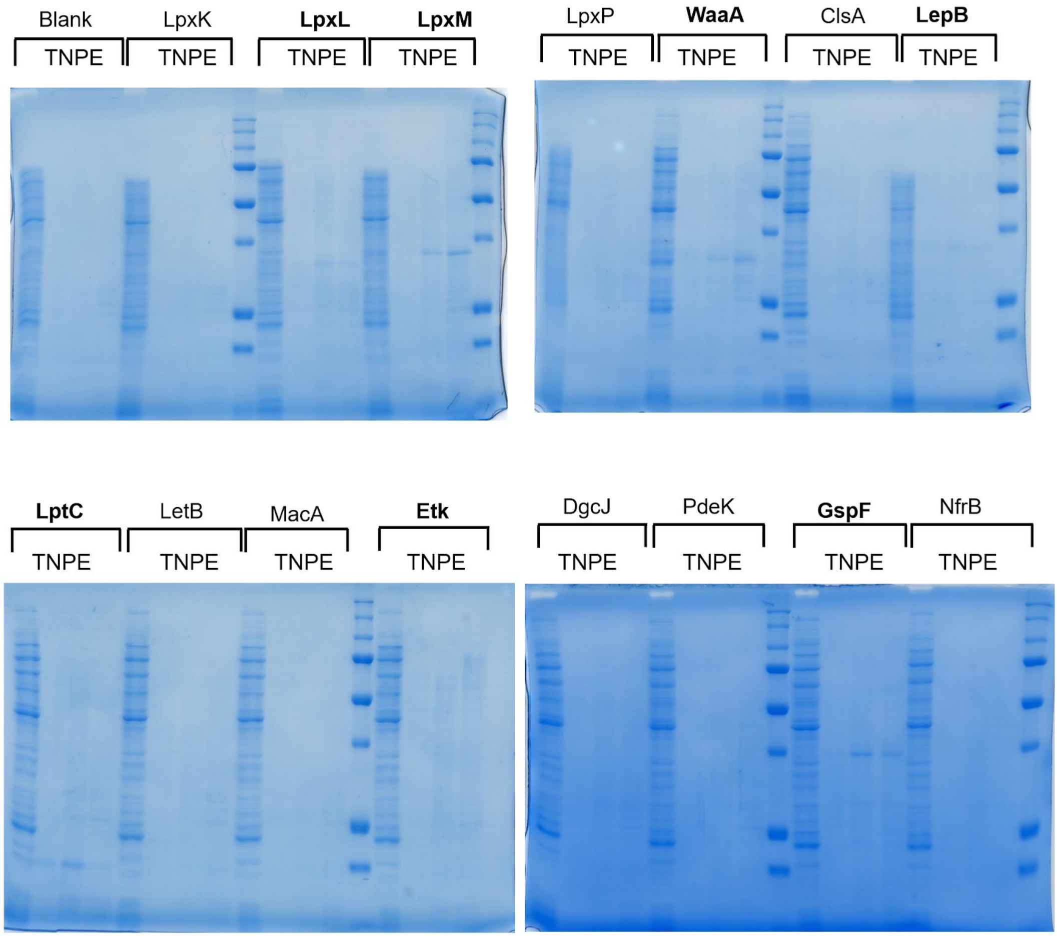


**Supplementary Figure 16**. SDS-PAGE Gels Group 8.

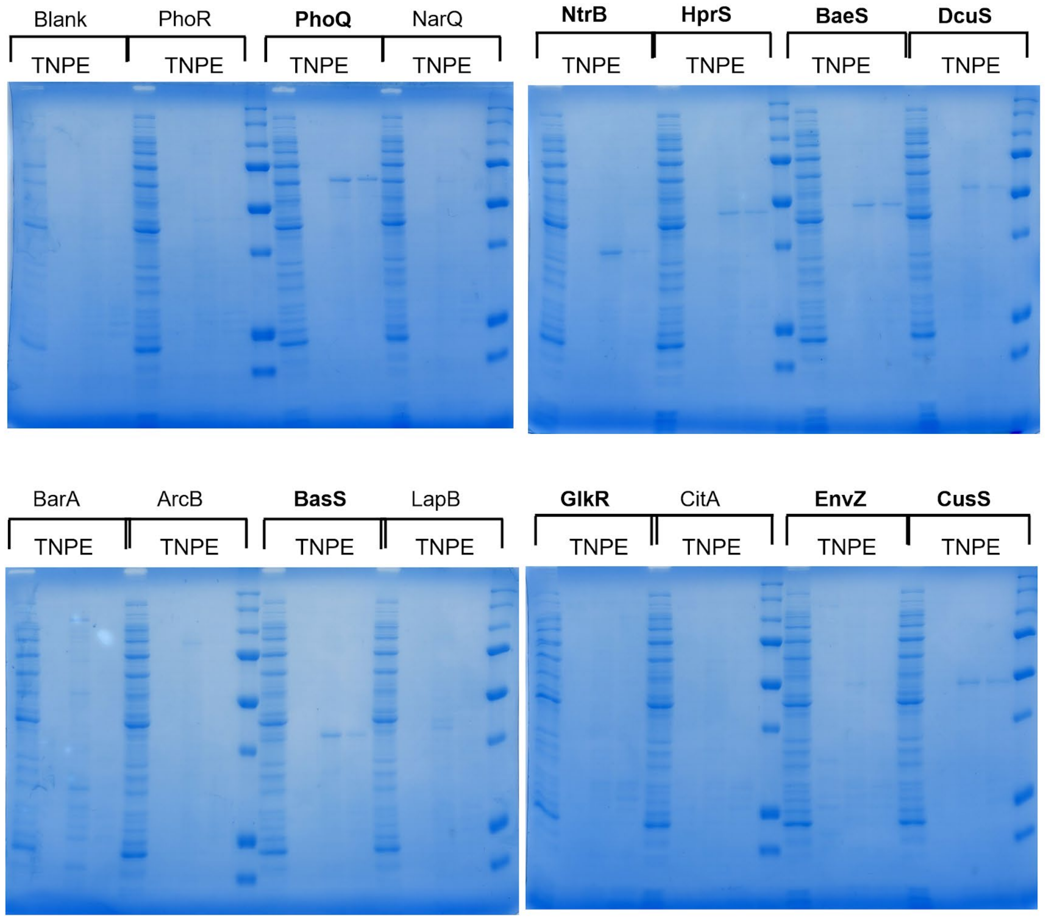


**Supplementary Figure 17**. SDS-PAGE Gels Group 9.

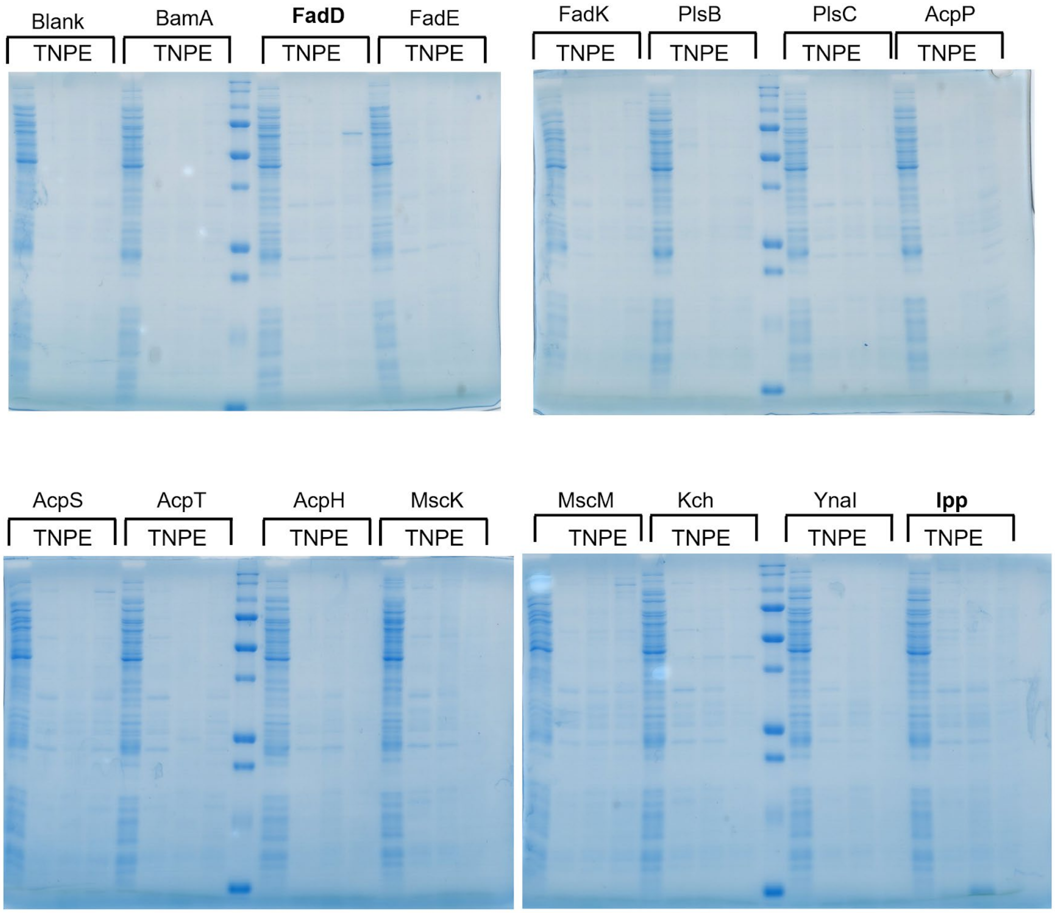


**Supplementary Figure 18**. SDS-PAGE Gels Group 10.

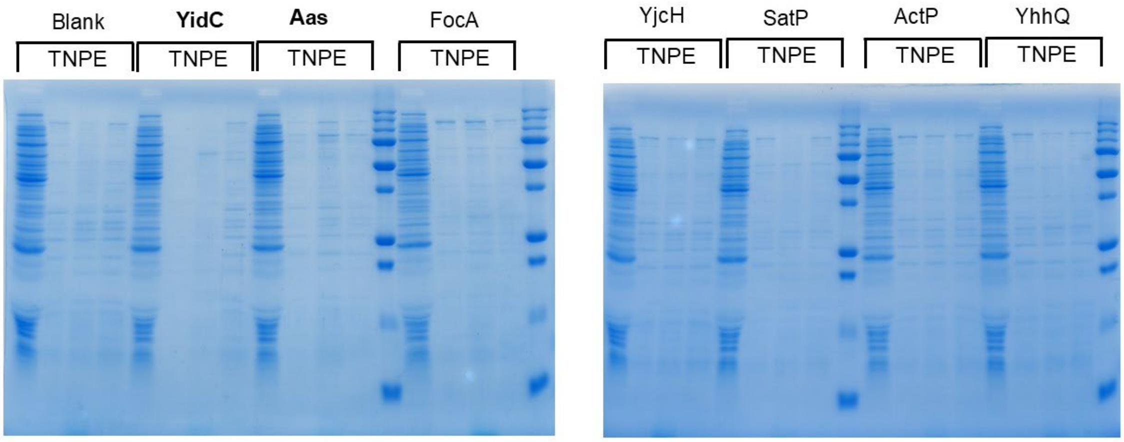


**Supplementary Figure 19**. SDS-PAGE Gels Group 11.

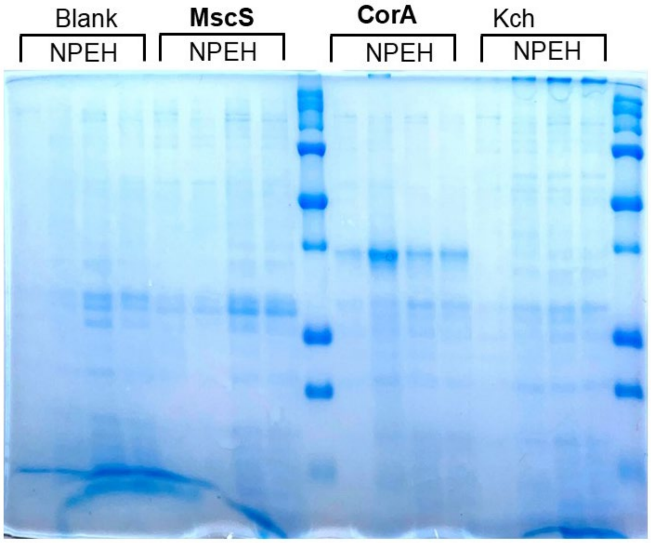


**Supplementary Figure 20.** Gel Group 12. SDS-PAGE for MscS, CorA, and Kch. The lanes are 'No SLB', 'DOPC SLB', 'ECL SLB', and 'Hybrid SLB'.

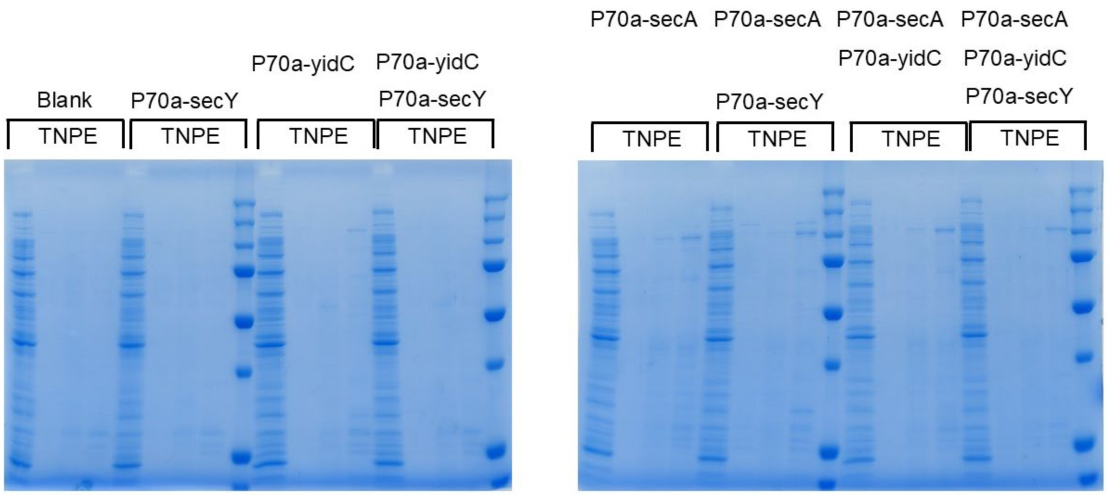


**Supplementary Figure 21.** SDS-PAGE for combinations of P70a-*secY,* P70a-*yidC*, and P70a-*secA*.

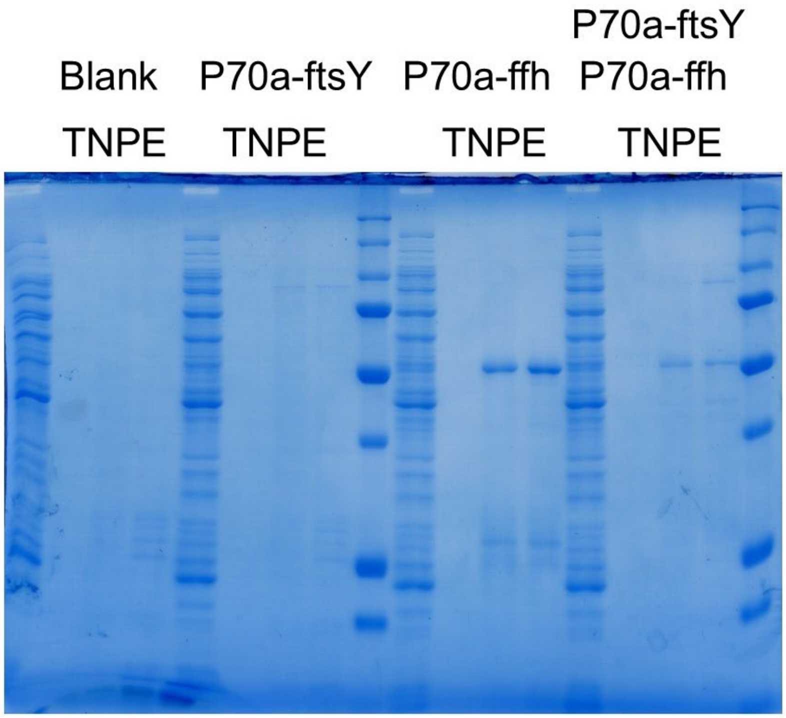


**Supplementary Figure 22.** SDS-PAGE for combinations of P70a-*ftsY* and P70a-*ffh*.

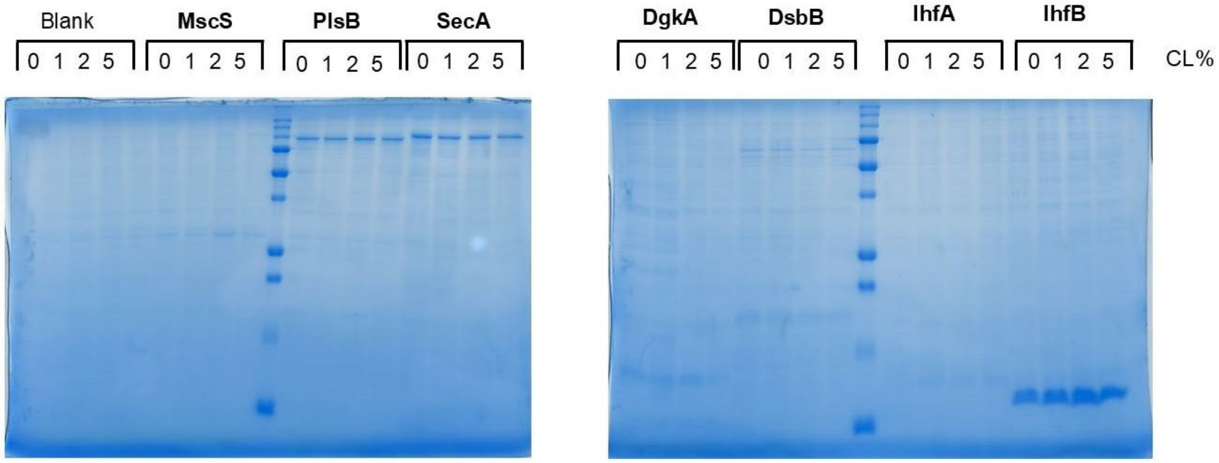


**Supplementary Figure 23**. Effect of CL on the binding for a selection of proteins.

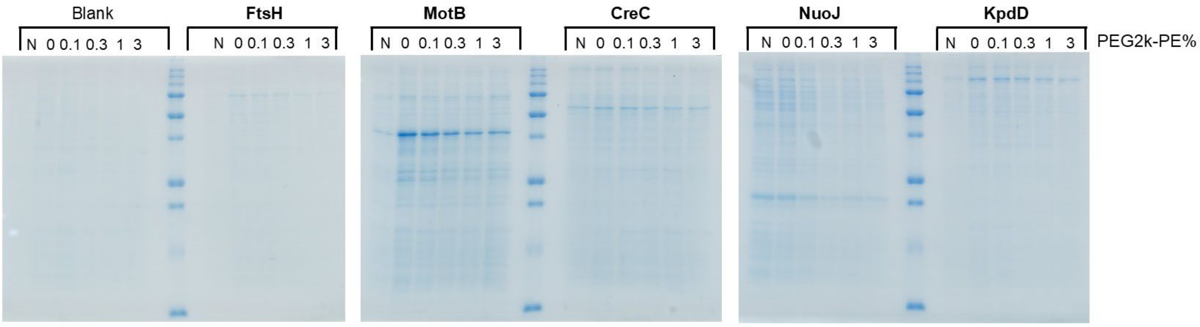


**Supplementary Figure 24**. Effect of PEG2k-PE on the binding for a selection of proteins. PEG2k-PE is added to ECL SLBs.

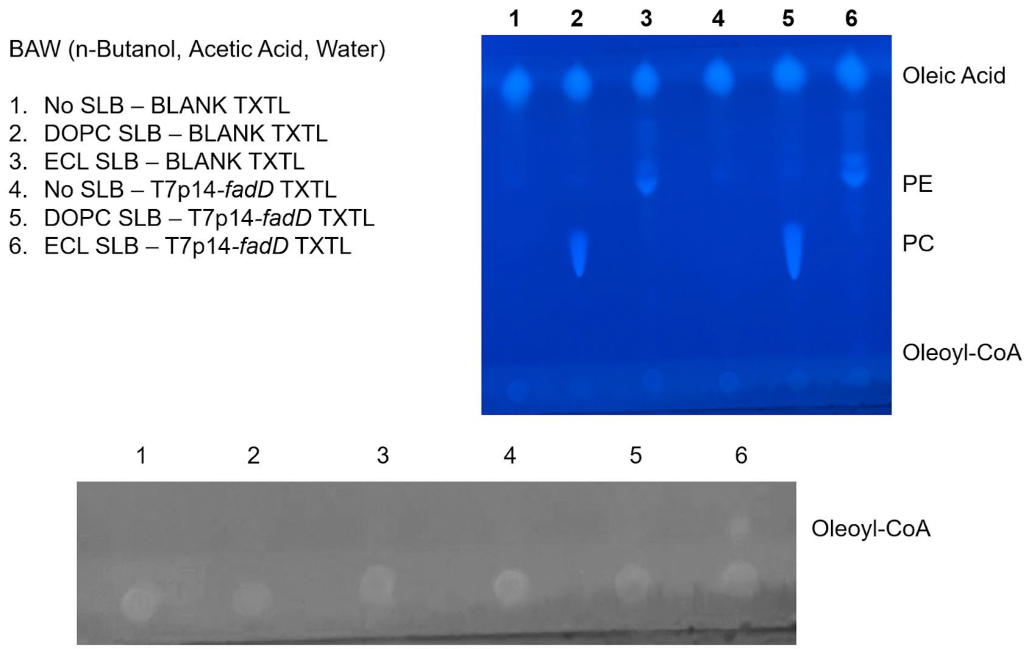


**Supplementary Figure 25.** Visualization of FadD activity with Primulin staining.

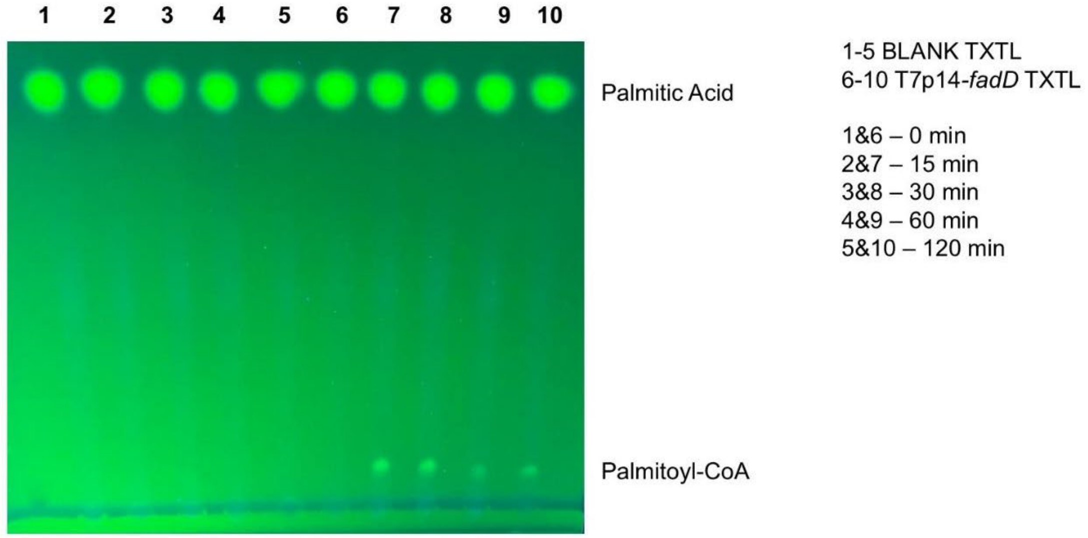


**Supplementary Figure 26.** Time course of palmitoyl-CoA synthesis.

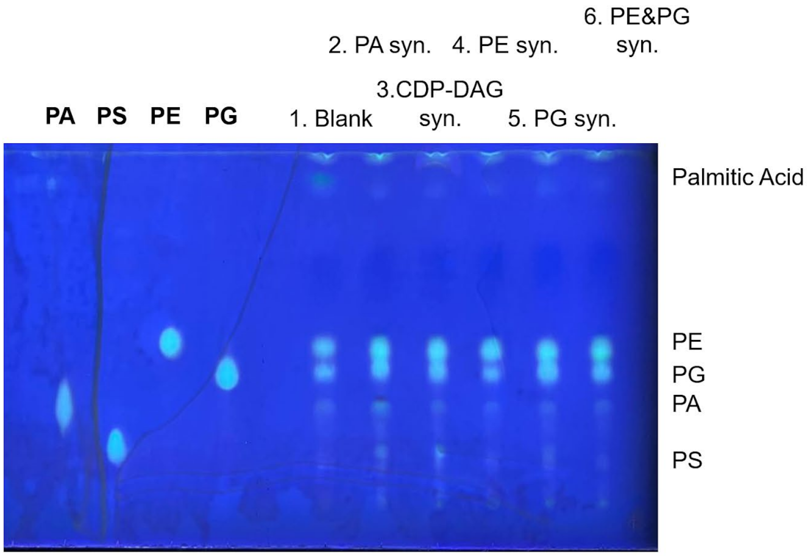


**Supplementary Figure 27.** Visualization of lipid synthesis combinations with Primulin staining.

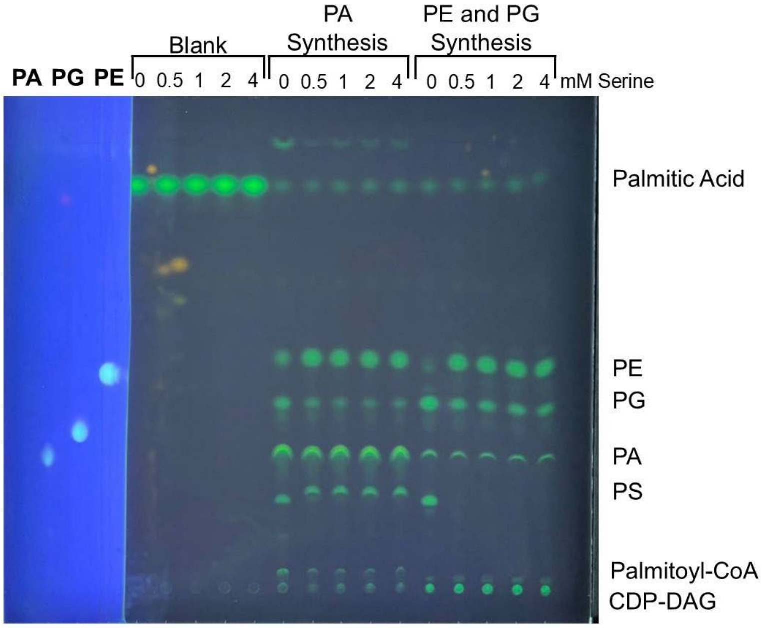


**Supplementary Figure 28.** L-Serine range for partial and full lipid synthesis pathways. The protein standards were dipped in primulin.
